# Deep mutational scanning of a multi-domain signaling protein reveals mechanisms of regulation and pathogenicity

**DOI:** 10.1101/2024.05.13.593907

**Authors:** Ziyuan Jiang, Anne E. van Vlimmeren, Deepti Karandur, Alyssa Semmelman, Neel H. Shah

**Affiliations:** Department of Chemistry, Columbia University, New York, NY 10027; Department of Biological Sciences, Columbia University, New York, NY 10027; Department of Biochemistry, Vanderbilt University, Nashville, TN 37232

**Keywords:** SHP2, *PTPN11*, tyrosine phosphatase, deep mutational scanning, allostery, molecular dynamics

## Abstract

Multi-domain signaling enzymes are often regulated through extensive inter-domain interactions, and disruption of inter-domain interfaces by mutations can lead to aberrant signaling and diseases. For example, the tyrosine phosphatase SHP2 contains two phosphotyrosine recognition domains that auto-inhibit its catalytic domain. SHP2 is canonically activated by binding of these non-catalytic domains to phosphoproteins, which destabilizes the auto-inhibited state, but numerous mutations at the main auto-inhibitory interface have been shown to hyperactivate SHP2 in cancers and developmental disorders. Hundreds of clinically observed mutations in SHP2 have not been characterized, but their locations suggest alternative modes of dysregulation. We performed deep mutational scanning on full-length SHP2 and the isolated phosphatase domain to dissect mechanisms of SHP2 dysregulation. Our analysis revealed mechanistically diverse mutational effects and identified key intra- and inter-domain interactions that contribute to SHP2 activity, dynamics, and regulation. Our datasets also provide insights into the potential pathogenicity of previously uncharacterized clinical variants.

## Introduction

The majority of human signaling proteins contain multiple domains with distinct biochemical functions. These domains facilitate catalytic activity, such as phosphorylation and dephosphorylation, or binding to ligands, including short linear protein motifs^1^, phosphorylated protein residues^2^, lipids^3^, and nucleic acids^4^. Regulatory interactions between these functional domains and inter-domain linkers enable multi-domain signaling proteins to function as highly regulatable switches capable of accessing different conformations in response to specific signals^5,6,7^. While this metastability is critical for regulation, it also makes multi-domain signaling proteins highly susceptible to dysregulation by mutations, which can disrupt their intra-domain allostery, inter-domain interactions, or linker structure and dynamics, leading to aberrant signaling and diseases^8^. A quintessential example is the protein tyrosine phosphatase (PTP) SHP2. This enzyme has two phosphotyrosine recognition (SH2) domains N-terminal to its catalytic PTP domain (**Figure 1A**). In signaling pathways such as the Ras/Erk and Jak/Stat pathways, SHP2 is recruited via its SH2 domains to phosphotyrosine-bearing sequences on receptors and scaffold proteins (**Figure 1B**). Upon phosphoprotein binding, SHP2 switches from a closed, auto-inhibited state characterized by extensive N-SH2/PTP domain interactions, to an open, active state, allowing for the dephosphorylation of downstream substrates (**Figure 1B,C**)^9^.

**Figure 1.**
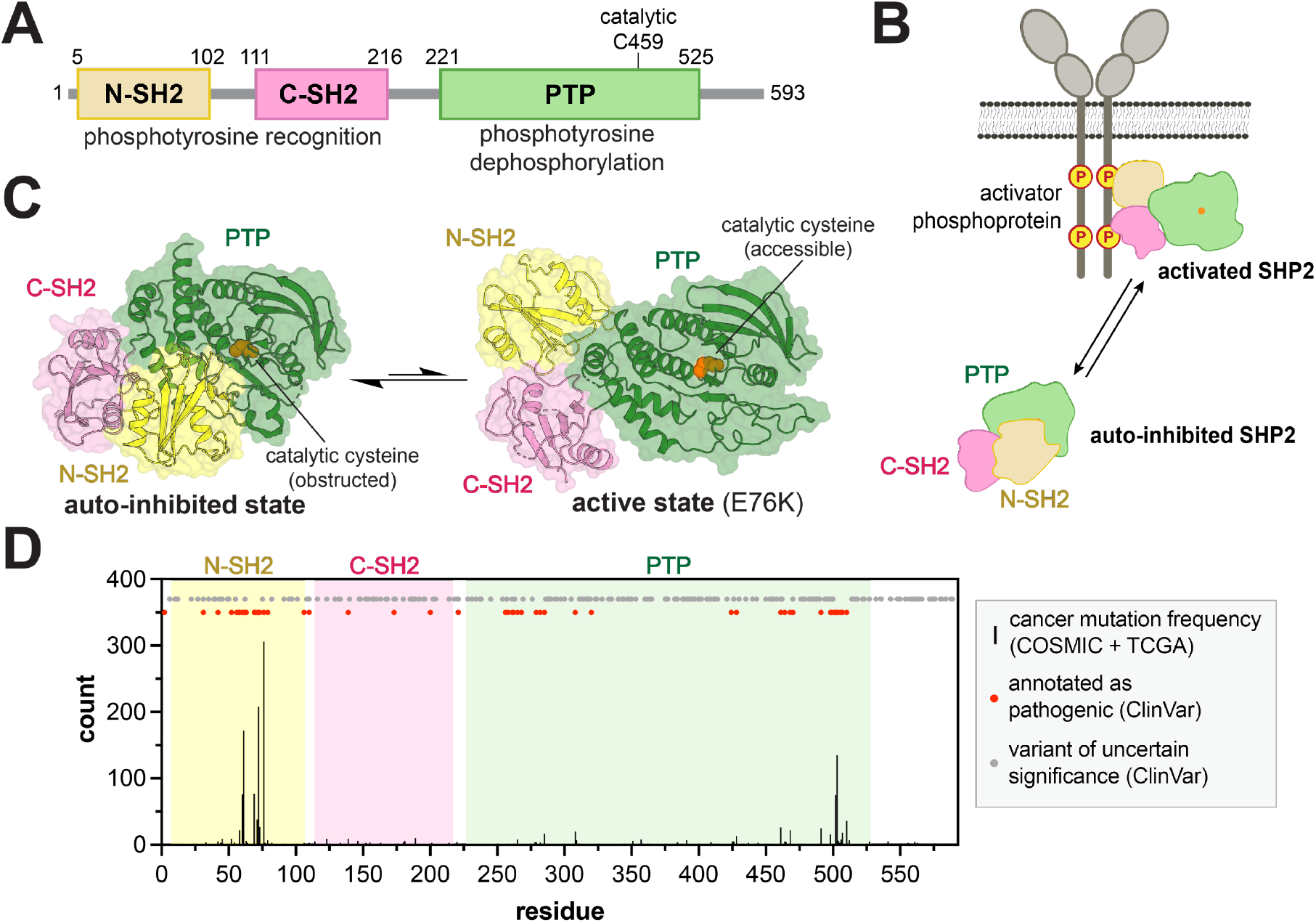
SHP2 activation and dysregulation by mutations. (**A**) Domain architecture diagram of SHP2. (**B**) Cartoon depiction of SHP2 activation by phosphoprotein binding. (**C**) Structures of the auto-inhibited (PDB code 4DGP) and active states of SHP2 (PDB code 6CRF). Note that the active state shown is that of the E76K mutant, and other active conformations likely exist in solution. (**D**) Positions and frequencies of cancer hotspot mutations (black bars) in SHP2 along its 593-residue sequence, derived from the COSMIC and TCGA databases. Sites of other pathogenic mutations and variants of uncertain significance are labeled as red and gray dots, respectively, derived from the ClinVar database.

Missense mutations in the SHP2 coding gene (*PTPN11*) can cause cancers and developmental disorders^10^. The functional effects of mutations in SHP2 vary across disease contexts. For example, in the developmental disorders Noonan Syndrome and Noonan Syndrome with Multiple Lentigines, gain-of-function SHP2 mutants confer their pathogenicity by promoting Ras/Erk activation, but some loss-of-function mutants can have similar phenotypic effects^11^. Numerous cases of gain-of-function SHP2 mutants are found in hematopoietic cancers^12,13^, and SHP2 also plays critical roles in the development of different types of solid tumors^14–16^. However, SHP2 has also been demonstrated to function as a tumor suppressor in liver cancer and a few other cancer types^17–19^. The spectrum of phenotypic effects caused by SHP2 variants is likely reflective of different axes of SHP2 function, including intrinsic catalytic activity, allosteric regulation, and protein-protein interactions. Given this, rigorous biochemical and biophysical characterization of disease-associated SHP2 mutants has substantially informed our understanding of SHP2 dysregulation and pathogenicity. For example, the extensive characterization of activating SHP2 mutants, including the high-frequency cancer mutation E76K, revealed key features of the N-SH2/PTP auto-inhibitory interface and also yielded the first active-state structure of SHP2^20,21^. The Noonan Syndrome T42A mutation in the N-SH2 domain was found to alter SHP2 ligand affinity and specificity, sensitizing SHP2 to activators^22,23^. The Y279C mutation, found in Noonan Syndrome with Multiple Lentigines, has been shown to disrupt phosphoprotein binding in the PTP active site, thereby significantly diminishing catalytic activity^24^. However, hundreds of disease-associated SHP2 mutants remain poorly studied^25^. These mutants span all three domains of SHP2 and the linkers between them, and many of them lack obvious mechanistic explanations of pathogenicity, making their significance unclear (**Figure 1D**). Characterization of these mutations will help elucidate novel features of SHP2 structure, regulation, and pathogenicity.

Deep mutational scanning is a powerful method for characterizing protein mutants^26^. By combining selection assays on pooled libraries with deep sequencing, this method provides a way to profile mutational effects across a protein with high throughput. Deep mutational scanning has been applied to reveal structure-function relationships^27^, predict protein structures and dynamics^28,29^, map drug resistance^30^, and examine stability and expression^31,32^. Molecular insights into structure and regulation derived from deep mutational scans have proven to be useful to rationalize pathogenicity^33–35^. Here, we present a deep mutational scanning platform to characterize the effects of SHP2 mutations on phosphatase activity. We used this platform to examine comprehensive point mutant libraries of both full-length SHP2 (SHP2_FL_) and its isolated phosphatase domain (SHP2_PTP_). These experiments yielded the activity profile of over 11,000 SHP2 mutants, including previously uncharacterized variants documented in disease databases. A comparative analysis of the two datasets, supported by molecular dynamics (MD) simulations, revealed several classes of mutations with distinct mechanisms of dysregulation and pathogenicity. Our analysis also identified novel interactions that mediate the transition of SHP2 between its auto-inhibited and active states, providing new insights into SHP2 regulation.

## Results

### Rescue of yeast growth from tyrosine kinase toxicity enables SHP2 deep mutational scanning

We developed a yeast viability assay in which cell growth is dependent on the catalytic activity of SHP2. Lacking significant endogenous tyrosine kinase/phosphatase signaling, yeast (*S. cerevisiae*) proliferation is arrested when expressing an active tyrosine kinase, whereas co-expression of an active tyrosine phosphatase can rescue yeast growth^36–38^. We co-expressed SHP2_FL_ variants known to have different levels of catalytic activity with two active versions of Src kinase, full-length viral Src (v-Src_FL_) and the isolated c-Src kinase domain (c-Src_KD_). In the presence of either kinase, the rate of yeast growth was dependent on the catalytic activity of the SHP2 variant (**Supplementary Figure 1A**). Moreover, the activity of the tyrosine kinase dictates the selection pressure of the assay. With the highly active v-Src_FL_ kinase, the more active SHP2 variants were better differentiated, while with the less active c-Src_KD_ kinase, lower activity SHP2 variants were easier to differentiate (**Supplementary Figure 1B,C**).

To comprehensively characterize SHP2 mutant activity, we conducted selection assays with SHP2_FL_ and SHP2_PTP_ saturation mutagenesis libraries co-expressed with each Src kinase variant (**Figure 2A**). Libraries were constructed using the Mutagenesis by Integrated TilEs (MITE) method, with SHP2_FL_ and SHP2_PTP_ divided into 15 and 7 sub-libraries (tiles), respectively (**Supplementary Table 1**)^39^. Each sub-library was separately introduced into yeast cells alongside plasmids encoding either v-Src_FL_ or c-Src_KD_. Cells were then subject to selection by induction of kinase and phosphatase expression, followed by a 24-hour outgrowth phase. Before and after outgrowth, the SHP2-coding DNA was isolated and deep sequenced, allowing for the calculation of enrichment scores for each variant, relative to wild-type SHP2 (**see Methods**). Two to four replicates of high-quality selection data were acquired for each sub-library (each tile), with good correlation of enrichment scores between replicates (**Supplementary Figure 2A**). The average enrichment scores of all variants across replicates were plotted as heatmaps. Due to the selection pressures of the two conditions and differences in intrinsic catalytic activity between SHP2_FL_ and SHP2_PTP_, we observed that the SHP2_FL_+c-Src_KD_ and SHP2_PTP_+v-Src_FL_ selection assays provided the best dynamic range for both gain-and loss-of-activity mutants, and these datasets were used for subsequent analysis (**Figure 2B,C, Supplementary Figure 2B-D, and Supplementary Tables 2,3**).

**Figure 2.**
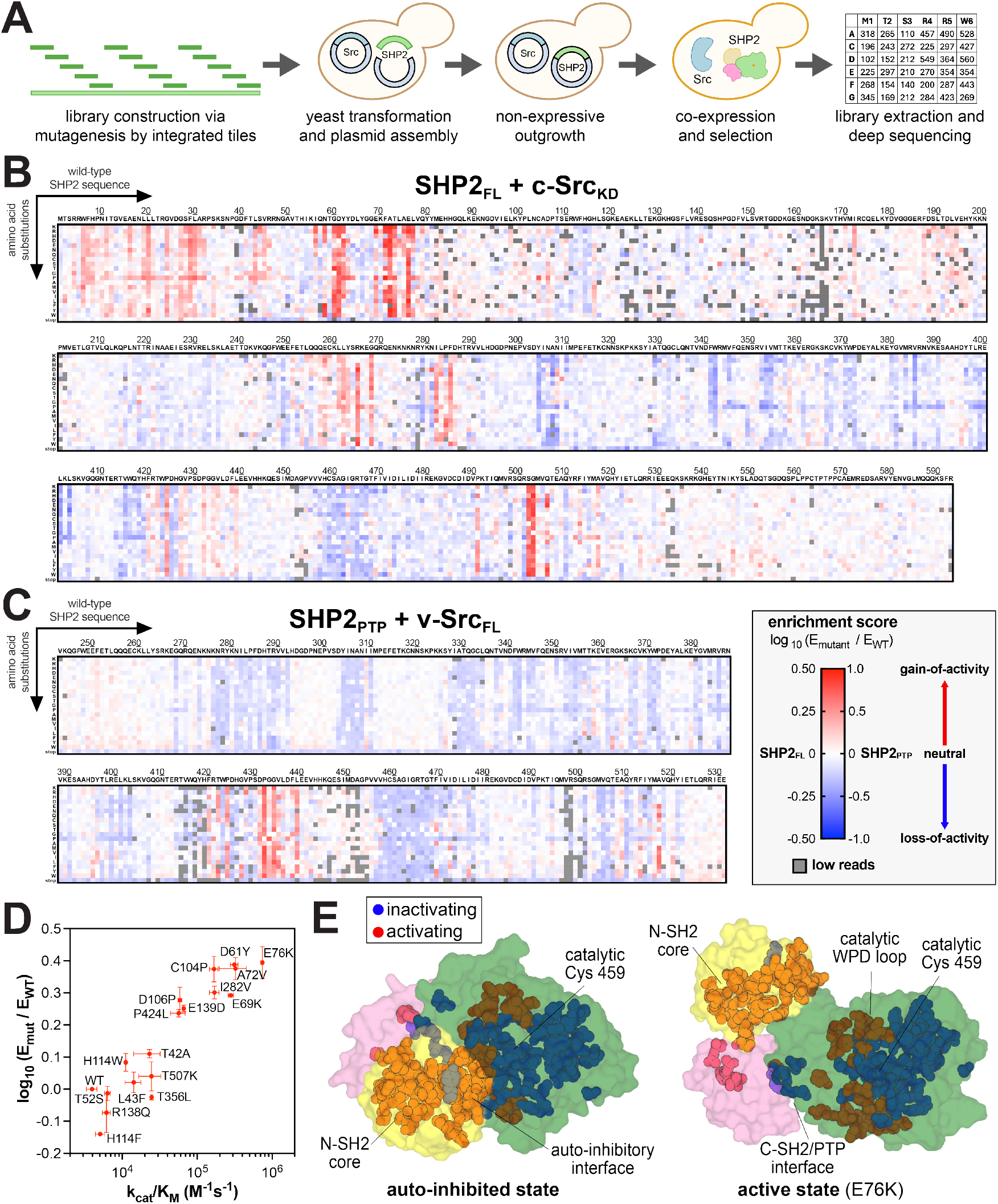
Deep mutational scanning of SHP2. (**A**) Workflow for SHP2 mutational scanning. SHP2 variant libraries were constructed using MITE, integrated into a yeast plasmid, co-expressed with Src, and subject to selection and deep sequencing. (**B**) Heatmap depicting the enrichment scores for SHP2_FL_ co-expressed with c-Src_KD_ (n = 2-4). Heatmap depicting the enrichment scores for SHP2_PTP_ co-expressed with v-Src_FL_ (n = 2). (**D**) Correlation between SHP2 variant enrichment scores in the SHP2_FL_ + c-Src_KD_ selection assay and measured catalytic activity against DiFMUP (n = 3). (**E**) Sites in the SHP2_FL_ mutational scan that are substantially activating (red) and inactivating (blue) on average, mapped on the auto-inhibited and active state structures of SHP2 (PDB codes 4DGP and 6CRF, respectively). Full mutational scanning datasets can be found in **Supplementary Tables 2 and 3**.

To validate that our selection assay faithfully reports on SHP2 catalytic activity as a function of mutations, we purified several full-length SHP2 mutants and measured their basal catalytic activities. The catalytic efficiencies (*k*_cat_/*K*_M_) of these mutants correlated well with their enrichment scores in the SHP2_FL_ selection (**Figure 2D and Supplementary Table 4**). Thus, while our mutational scans may be impacted by mutant-specific changes in expression level, protein-protein interactions, or substrate specificity, our results suggest that basal catalytic activity is the major determinant of enrichment in our selection assays. For further validation, we compared our datasets with known mutational effects. As expected, well-studied activating mutations at the N-SH2/PTP interface (e.g. E76, D61, and S502 substitutions) were highly enriched in the SHP2_FL_ selections (**Figure 2B,E**). Furthermore, mutations at key catalytic residues (e.g. C459 and D425) were depleted in both the SHP2_FL_ and SHP2_PTP_ selections (**Figure 2B,C**). In addition to the known mutations, we observed strong mutational hotspots in unexpected regions, including activating mutations in the core of the N-SH2 domain, inactivating mutations at the C-SH2/PTP interface, and activating mutations around the key catalytic WPD loop, which will be discussed in later sections (**Figure 2E**).

### Mutational scans reveal disease-specific profiles of SHP2 activity

Of the ∼600 of clinically-observed SHP2 variants, only 20% are annotated as pathogenic. Thus, we used our SHP2_FL_ enrichment scores to gain insights into the functional effects of clinical variants (**Supplementary Table 5**). When compared to the full distribution of mutational effects, mutations annotated as pathogenic were more gain-of-function on average, however many reported pathogenic mutations did not enhance SHP2 activity (**Figure 3A,B**). This is consistent with the presence of some known loss-of-activity SHP2 mutations in developmental disorders^40^. High-frequency cancer mutations skewed further toward gain-of-activity, but even in this category, a few were neutral or even loss-of-activity (**Figure 3B**). It is noteworthy that many low-frequency cancer mutations, reported in fewer than five cases across COSMIC and TCGA, were neutral or loss-of-activity in the SHP2_FL_ selection assay. Most of these mutations have not been characterized, but they may still drive oncogenic signaling through mechanisms that do not rely on phosphatase activity, such as scaffolding mediated by the SH2 domains.

**Figure 3.**
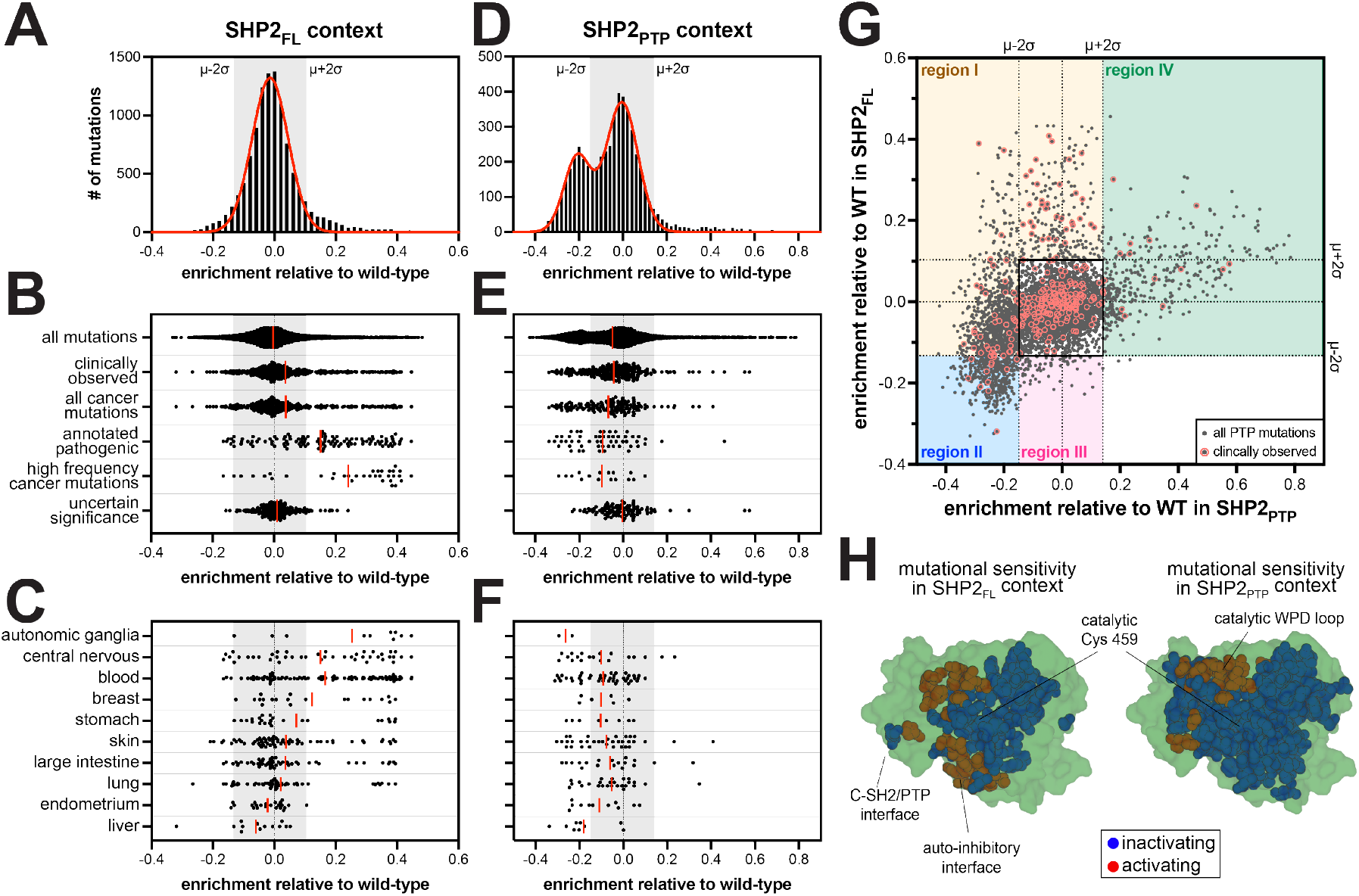
Differential phosphatase domain mutational effects in SHP2_FL_ and SHP2_PTP_ contexts. (**A**) Full distribution of enrichment scores from the SHP2_FL_ + c-Src_KD_ selection, fit to a single Gaussian curve. (**B**) Distribution of enrichment scores from the SHP2_FL_ + c-Src_KD_ selection for various sets of clinically relevant mutants. (**C**) Distribution of enrichment scores from the SHP2_FL_ + c-Src_KD_ selection for mutants found in different cancer subtypes. (**D**) Full distribution of enrichment scores from the SHP2_PTP_ + v-Src_FL_ selection, fit to the sum of two Gaussian curves. (**E**) Distribution of enrichment scores from the SHP2_PTP_ + v-Src_FL_ selection for various sets of clinically relevant PTP domain mutants. (**F**) Distribution of enrichment scores from the SHP2_PTP_ + v-Src_FL_ selection for PTP domain mutants found in different cancer subtypes. Red lines in panels B, C, E, and F correspond to mean values. (**G**) Correlation between enrichment scores for PTP domain mutants in the SHP2_FL_ + c-Src_KD_ (y-axis) and SHP2_PTP_ + v-Src_FL_ (x-axis) selections. Mechanistically distinct regions are highlighted as different-colored boxes. The uncolored black box near the center of the graph encompasses mutations that have small or neutral functional effects in both selections. Clinically observed mutants are circled in red. (**H**) Sites in the SHP2_FL_ (*left*) and SHP2_PTP_ (*right*) mutational scans that are substantially activating (red) and inactivating (blue) on average, mapped on a structures of SHP2 PTP domain (PDB code 3ZM0).

We further parsed cancer mutations by the primary tissue type and were surprised to find differences in the distribution of SHP2 activity across cancer subtypes (**Figure 3C**). As expected, many blood cancer mutations were highly activating in the SHP2_FL_ mutational scan^11,41^. Gain-of-activity mutants were less common in other cancer types, but those observed were largely the same as the hallmark activating mutations observed in blood cancers (e.g. D61, A72, E76, and S502 substitutions). The prevalence of neutral and loss-of-function mutants in many solid tumor types raises the question of whether those mutations are drivers of pathogenic signaling or simply bystanders. A striking observation from our analysis that supports a pathogenic role for loss-of-function SHP2 mutations is that all documented SHP2 liver cancer mutants were neutral or loss-of-activity in our dataset (**Figure 3C and Supplementary Table 5**). SHP2 is known to be a tumor suppressor in liver cancer, but this has only been established by knock-out and knock-down models, not missense mutations^17^. Given that all liver cancer mutants were neutral or loss-of-activity, we hypothesize that mutants in this context are pathogenic through loss of SHP2 tumor suppressor activity. As such, these mutants may warrant further investigation using *in vivo* models.

### Impaired catalytic activity is compensated by disrupted auto-inhibition in many disease mutants

Many unstudied SHP2 disease mutants are found in the PTP domain. Our SHP2_PTP_ selection assays show that the majority of these disease mutants reduce the intrinsic phosphatase activity of SHP2 (**Figure 3D-F**). To better understand the molecular basis for pathogenicity of PTP domain mutations, we juxtaposed enrichment scores from SHP2_PTP_ and SHP2_FL_ selections (**Figure 3G**). A monotonic relationship between the two datasets would suggest that the signal in both datasets depends solely on phosphatase domain activity. However, the enrichment scores follow a funnel-like distribution, spanning all four quadrants of the two-dimensional space, allowing us to decouple mutational effects on intrinsic catalytic activity from those that alter inter-domain interactions (**Figure 3G**). To narrow our analysis, we focused on significantly activating or deactivating mutants in either selection that fall more than two standard deviations from the mean of the distribution centered around wild-type (**Figure 3A,D,G**). We binned these mutations into four distinct regions (**Figure 3G**, *colored boxes*). Disease mutations in each of the regions dysregulate SHP2 through distinct mechanisms (**Figure 3G**, *circled points*).

Region I encompasses mutations in the PTP domain that disrupt auto-inhibition but have either a neutral or negative effect on intrinsic phosphatase activity. Many well-studied mutations in this region exclusively disrupt the auto-inhibitory interface, with a nominal effect on SHP2_PTP_ activity (e.g. cancer mutations at R265 and S502, which interact with E76 in auto-inhibited SHP2). By contrast, our datasets reveal a number of previously unstudied disease mutants in region I that significantly reduce intrinsic phosphatase activity (depleted in SHP2_PTP_ dataset) but compensate for this perturbation by alleviating auto-inhibition (neutral or enriched in SHP2_FL_ dataset) (**Supplementary Table 5**). Many of them are positioned close to key catalytic loops that regulate both catalysis and auto-inhibition, such as D286, H287, T288, N306, and N308 mutants underneath the substrate-binding pTyr loop, or R501, M504, and V505 mutants in the Q loop region (**Supplementary Figure 3A,B**). Frequent disease mutations at P491 also fall into this category, likely by disrupting C-SH2/PTP interactions in the closed conformation to alleviate auto-inhibition (**Supplementary Figure 3B,C**). Previous studies have demonstrated that mutants with reduced auto-inhibition more readily interact with binding partners in signaling pathways, leading to increased signaling activity^22,24,42^. The prevalence of disease-associated mutants in region I with a substantial loss of intrinsic phosphatase activity supports the idea that there are important roles of SHP2 in disease contexts that depend on protein-protein interactions but not protein dephosphorylation^43^.

More dramatic decreases in PTP domain activity cannot be fully compensated by reduced auto-inhibition, as exemplified by previously studied region I mutants T468M and T507K^42,44^. For these mutants, where SHP2_PTP_ activity is significantly impaired, disrupted auto-inhibition in SHP2_FL_ can only bring SHP2_FL_ activity back to wild-type levels (**Supplementary Figure 3A**). Many region II mutants represent extreme cases where reduced PTP activity cannot be overcome by relieved auto-inhibition. For example, the disease-associated mutant Y279C at the region I/II boundary disrupts auto-inhibition, but it has such intrinsically low catalytic activity that it is still significantly depleted in the SHP2_FL_ selections (**Supplementary Figure 3A**)^24,45^. To corroborate this idea, we measured melting temperatures of several full-length mutants by differential scanning fluorimetry, which has been shown to report on SHP2 auto-inhibition^46^. Indeed, Y279C, T468M, and T507K have lowered melting temperatures compared to wild-type, indicating a less auto-inhibited conformation (**Supplementary Figure 3D**). Several other clinically-observed mutants in region II lie in the PTP active site and at the auto-inhibitory interface, including unstudied mutants T357M and I463L (**Supplementary Figure 3A,C**). By analogy to well-characterized region I and II mutations, we hypothesize that these uncharacterized loss-of-function mutants disrupt auto-inhibition and contribute to pathogenicity via phosphatase-activity-independent signaling roles.

### Residues at the C-SH2/PTP interface mediate exit from the auto-inhibited state of SHP2

Region III mutations are deactivating in the SHP2_FL_ context but neutral in SHP2_PTP_. Many of them, most notably at E249, cluster on the α-helices facing the C-SH2 domain, and interactions at this interface differ between the auto-inhibited and active states (**Supplementary Figure 4A,B**). We hypothesize that this interface mediates the transition from auto-inhibited to open conformations, even in the absence of any activating ligands. Two key residues on the SH2 side of this interface are R111 and H114. Most mutations at these residues deactivate SHP2_FL_, and both residues harbor mutations reported in cancer (**Figure 4A and Supplementary Table 5**). The interaction between R111 and E249 has been reported to stabilize an active conformation of SHP2^47^, however the roles of these residues in SHP2 dynamics are unexplored, and H114 has not been implicated in SHP2 regulation.

**Figure 4.**
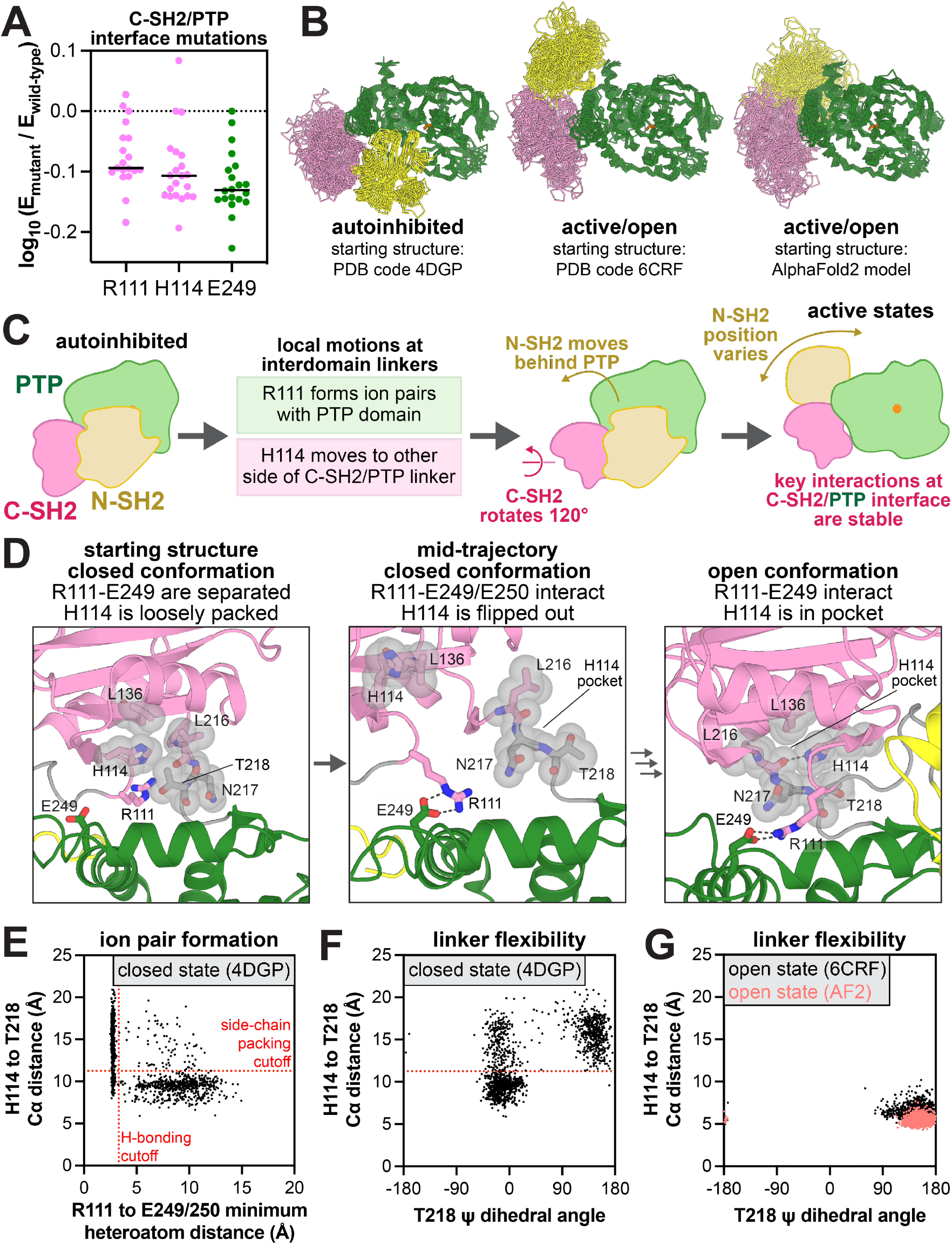
Structure and dynamics at the C-SH2/PTP interface. (**A**) Mutational effect at R111, H114, and E249 in the SHP2_FL_ selection with c-Src_KD_. (**B**) Conformational sampling across 18 MD simulations of SHP2 starting from three different conformational states. (**C**) Hypothesized sequence of events in the SHP2 closed-to-open transition. (**D**) Representative frames from MD simulations of SHP2 highlighting a lack of R111-E249 ion pairing and loose packing of H114 in simulations starting from the closed conformation (*left*), formation of the R111-E249 ion pair with concomitant exit of H114 from the C-SH2/PTP linker pocket in simulations starting from the closed conformation (*middle*), and stable R111-E249 and H114-L216 interactions in simulations starting from the open conformation (*right*). The transition from the middle to right frame is not observed in our simulations. (**E**) Correlation between R111-E249/E250 distance (shortest distance between one of the arginine terminal nitrogens and one of the four glutamate carbonyl oxygens) and H114-T218 Cα distance in the SHP2 closed conformation simulations. (**F**) Correlation between T218 ψ dihedral angle and H114-T218 Cα distance and in the SHP2 closed conformation simulations. (**G**) Correlation between R111-E249/E250 distance and H114-T218 Cα distance in the SHP2 open conformation simulations.

To examine how the C-SH2/PTP interface governs SHP2 regulation, we performed molecular dynamics (MD) simulations on near-full-length SHP2 (excluding the disordered tail), starting from three different conformational states: the auto-inhibited state (PDB code 4DGP), a crystallographic open state of the E76K mutant (PDB code 6CRF), and an alternative open state from an AlphaFold2 model, which has the N-SH2 domain further behind the PTP domain when compared to the crystal structure (**Figure 4B**). We used two open structures given the evidence that SHP2 adopts multiple active conformational states^20,48,49^. Starting from each state, we conducted three 2.5 μs simulations with both the wild-type sequence and the E76K mutant. In the open conformation simulations, starting from both the crystal structure and the AlphaFold2 model, the C-SH2/PTP interface is highly similar and quite stable. R111 frequently forms ion pairs with E249 and E250, or occasionally with E232 (**Supplementary Figure 4C,D**). Furthermore, H114 docks in a pocket consisting of L136, L216, N217, and T218, stabilized by hydrophobic interactions as well as a hydrogen bond between its imidazole nitrogen and L216 main chain carbonyl (**Figure 4D**, *right panel*, **Supplementary Figure 4E**). Our mutagenesis data show that aromatic residues with hydrogen bonding capacity (H and W) stabilize this active state, whereas other residues, including F, reduce SHP2 activity (**Supplementary Figure 4F and Supplementary Table 4**).

The C-SH2/PTP interface is more dynamic in the closed conformation simulations, and motions around R111 and H114 suggest a plausible sequence of events for the initiation of SHP2 activation (**Figure 4C,D**). In contrast to the open conformation, R111 does not participate in significant interactions in the closed conformation starting structure and points away from the PTP domain, while H114 engages T218 through hydrophobic interactions with its side chain (**Figure 4D**, *left vs right panels*). During the simulations, R111 rotates into the C-SH2/PTP cleft and intermittently forms ion pairs with E249 or E250, despite the lack of large inter-domain rearrangements (**Figure 4D**, *middle panel*). When R111 interacts with E249/E250, this movement disrupts the interaction between H114 and T218 (**Figure 4D**, *middle panel*, **Figure 4E**). Without this stabilizing interaction, the C-SH2/PTP linker becomes dynamic, enabling T218 to adopt multiple conformations (**Figure 4F**). This allows the C-SH2 domain to rotate around the C-SH2/PTP interface and sample the tighter stabilized conformation in the SHP2 open structures (**Figure 4D**, *right panel*, **Figure 4G**). Our mutational scanning data support the notion that T218-H114 packing must be disrupted for activation: substitution of T218 with smaller residues weakens the interaction with H114 and increases SHP2 activity, while bulkier substitutions enhance the H114 interaction and decrease SHP2 activity (**Supplementary Figure 4G**). Full inter-domain rearrangement cannot be captured on our simulation timescale, but the extensive reorganization of interactions at the C-SH2/PTP interface between the auto-inhibited and active states, and the mutational effects of residues on the interface in our data suggest that the interface is intimately involved in SHP2 activation.

Based on our analysis, loss-of-activity mutations at this interface operate by stabilizing the closed conformation of SHP2. These mutations would make SHP2 less capable of binding upstream phosphoproteins, as demonstrated previously for E249^47^. Consequently, no disease-associated PTP mutations appear in region III (**Figure 3G**). On the other hand, the T218S and T218A mutations are observed in developmental disorders (**Supplementary Table 5**). The functional effects of these mutations were previously unknown, but our results show that disruption of the T218-H114 interaction in the auto-inhibited conformation promotes SHP2 activation (**Supplementary Figure 4G**). We also note that the R111M and H114Y mutations have been observed in skin cancer, suggesting that there could be a role for loss-of-function mutations at the C-SH2/PTP interface in human diseases.

### Activating SHP2_PTP_ disease mutants regulate SHP2-specific WPD loop motions

Gain-of-activity mutants in the SHP2_PTP_ construct reside within region IV, where their effects on the SHP2_FL_ construct are neutral or activating. While mildly-activating SHP2_PTP_ mutants are widely distributed, the most activating mutants cluster around the WPD loop (**Figure 5A and Supplementary Figure 5A**). WPD loop closure positions a key catalytic residue (D425) for catalysis. Thus, mutations in this cluster likely alter WPD loop dynamics and favor a WPD-closed conformation^50,51^. In SHP2_FL_, WPD loop closure is blocked in the auto-inhibited state by the steric hinderance from the N-SH2 domain (**Supplementary Figure 5B**). Consequently, the activating effects of some mutants in region IV are less significant in the SHP2_FL_ context when compared to those disrupting auto-inhibition (region I).

**Figure 5.**
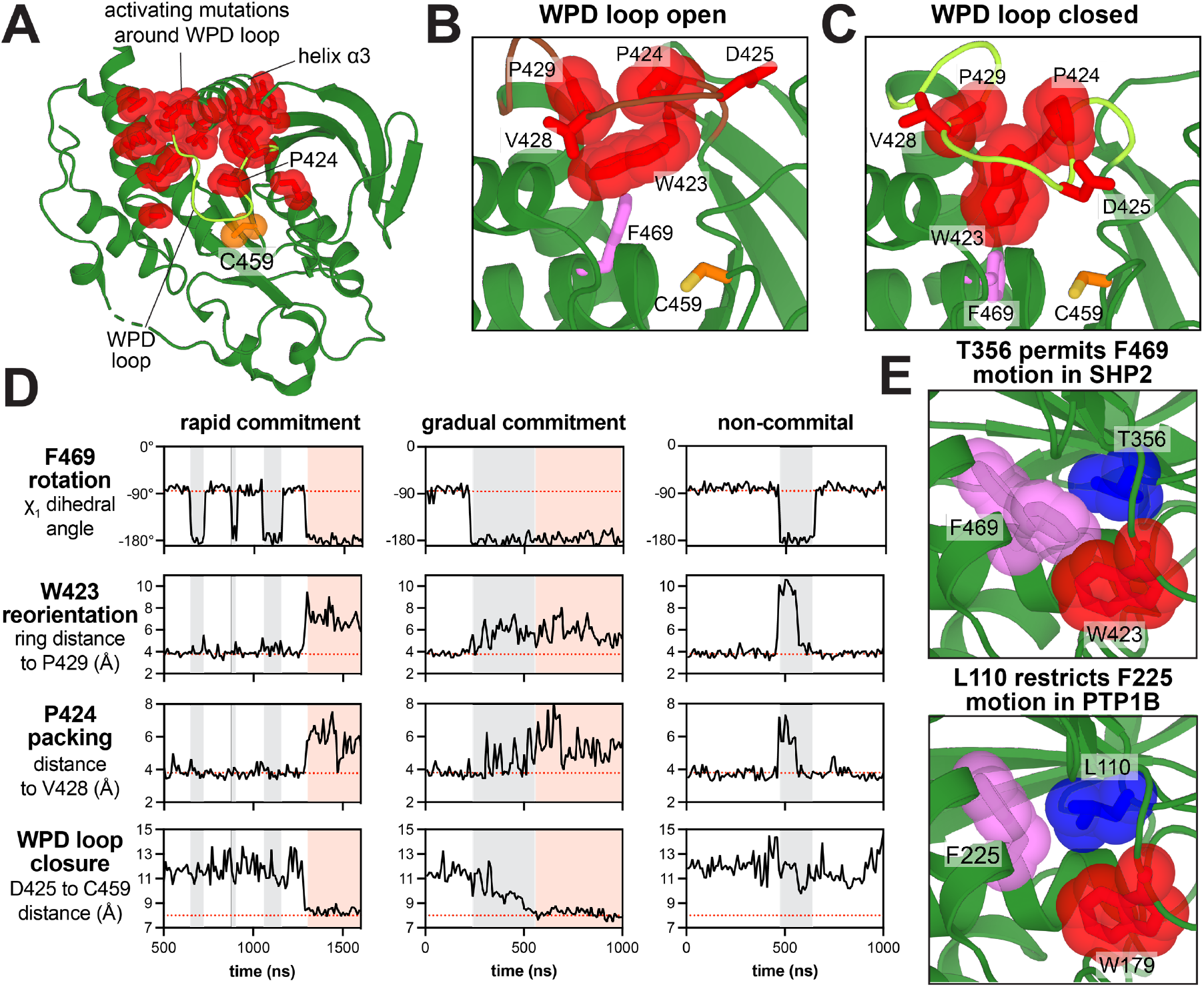
Mutational effects that alter WPD loop conformational dynamics. (**A**) Residues with consistent gain-of-function mutational effects around the WPD loop (PDB code 3ZM0). (**B**) Frame from MD trajectory of SHP2 starting from PDB 6CRF, showing packing of W423, P424, P429, and F469 in WPD loop open conformation. F469 points towards the WPD loop in this structure. (**C**) Frame from an MD trajectory starting from PDB 6CRF, showing that, in the WPD closed conformation, W423 displaces F469, and P424 is released from its intimate packing in the turn of the loop. (**D**) Three MD trajectories starting from the open conformation, showing coordinated movements between F469, W423, and P424. The first trajectory is an example of these movements culminating in WPD closure. The second trajectory also shows WPD loop closure, but more gradually. The third trajectory shows coupled F469, W423, and P424 motions that revert back to their starting state, resulting in no WPD loop closure. F469 rotation is quantified by χ_1_ angle of this residue. W423 reorientation is quantified by the distance between W423 Cζ_3_ and P429 Cγ. P424 packing is quantified by the distance between P424 Cγ and the V428 carbonyl. WPD loop closure is quantified by the distance between the D425 and C459 Cα atoms. Gray shaded segments denote F469 in a permissive state for WPD loop closure. Pink shaded segments denote WPD loop closure. (**E**) Small residue T356 (blue) permits F469 rotation in SHP2, restricting W423 rotation and WPD loop closure (PDB code 3ZM0, *top*). Multiple F469 rotamers are modeled in this crystal structure. Bulky residue L110 (blue) constrains F225 rotation in PTP1B, permitting facile W179 rotation and WPD loop closure (PDB code 5K9V, *bottom*).

In the MD trajectories of the active conformations of SHP2, we captured several instances of the WPD open-to-closed transition. These events reveal coordinated movements between key residues in this region and provide insights into how SHP2 may differ from other phosphatases, such as PTP1B (**Figure 5B-D**). When SHP2 adopts the WPD open conformation, W423, P424, and P429 pack together, and this cluster is buttressed by F469 (**Figure 5B**). Notably, F469 adopts multiple rotameric states in crystal structures and in our MD trajectories (**Figure 5D,E**)^52^. In the transition to the WPD closed conformation, F469 rotates away from the WPD loop, and W423 moves away from the loop center into the cleft that is otherwise occupied by F469 (**Figure 5C,D**). This transition releases P424 from its packing interactions, allowing it to access more diverse conformations. We quantified these movements by measuring the distance between W423 Cζ_3_ and P429 Cγ, the χ_1_ angle of F469, and the distance between P424 Cγ and the V428 carbonyl. In our simulations, we observed instances of concerted W423/F469/P424 movements that ultimately culminated in WPD loop closure, as well as instances where these residues sampled an on-pathway state and then reverted back to their inactive conformation, thereby aborting WPD loop closure. Our analysis shows that the F469 rotation acts as a gatekeeper for W423 movement and WPD loop closure (**Figure 5D**).

Although similar WPD loop motion has been observed before in the well-studied phosphatase PTP1B, the phenylalanine in PTP1B (F225) corresponding to SHP2 F469 only adopts one rotameric state that permits facile WPD loop opening and closing^53^. F225 is locked in this orientation in by the adjacent L110, while in SHP2 this leucine is substituted by a less bulky threonine (T356), allowing for the gatekeeping function of F469 (**Figure 5E**). Indeed, the T356L mutation in SHP2 is activating in our mutational scans and in biochemical assays, suggesting that F469 is held away from the WPD loop in this mutant (**Supplementary Figure 5C and Supplementary Table 4**). Notably, in human classical PTPs, only SHP2 and MEG2/PTPN9 have this Phe-Thr combination, making the gatekeeping feature unique to these two phosphatases (**Supplementary Figure 5D**).

Disease-associated mutations in the vicinity of the WPD loop can have a range of outcomes on SHP2 function. Mutations in this area that activate SHP2_PTP_ should promote WPD loop closure. For mutations just outside of the WPD loop that allosterically modulate loop dynamics, this activating effect is overridden by N-SH2 binding to the PTP domain. Potentially pathogenic mutations at F420, P432, G433, V435, and the gatekeeper F469 fall into this category (**Supplementary Figure 5A**). F469S, found in colon cancer, should favor WPD loop closure, and indeed this mutation activates SHP2_PTP_ in our mutational scan (**Supplementary Figure 5E**). Such mutations would only unleash their effects on signaling when bound to an activator phosphoprotein that displaces the N-SH2 domain. On the other hand, strongly activating mutations directly on the WPD loop can enhance PTP domain activity and also disrupt auto-inhibition. For example, all P424 mutants are activating in both SHP2_PTP_ and SHP2_FL_ screens, including brain cancer and Noonan Syndrome mutants P424S and P424L. In PTP1B, mutations at this conserved proline stabilize a WPD-closed conformation^51^, which would displace the N-SH2 domain from the auto-inhibitory interface in SHP2_FL_ (**Supplementary Figure 5B**). Indeed, basal activity and melting curve measurements confirm that P424L activates SHP2 and destabilizes its auto-inhibited state (**Supplementary Table 4 and Supplementary Figure 3D**). This dual-activating effect makes clinically-observed mutants that are activating in both screens very likely to be pathogenic. Intriguingly, the WPD loop cancer mutation V428M lies in region I instead of region IV (**Supplementary Table 5**), indicating that mutation alters the WPD loop conformation in a way that decreases SHP2_PTP_ activity while still disrupting auto-inhibition in the SHP2_FL_ construct.

### Pathogenic N-SH2 core mutants balance domain destabilization and phosphoprotein binding

Outside of the PTP domain, the N-SH2 domain is well-known to host disease mutations, most of which disrupt the auto-inhibitory interface. In addition to these interface mutations, the SHP2_FL_ mutational scan revealed a large cluster of activating mutations in the hydrophobic core of N-SH2 domain (W6, F7, I11, A16, L20, F29, L30, L43 and V45) (**Figures 2D and 6A**). We hypothesize that mutations at these sites disrupt hydrophobic packing, thereby destabilizing the N-SH2 domain and weakening its auto-inhibitory binding to the PTP domain. Consistent with this hypothesis, polar mutations at these sites are more activating than nonpolar mutations, as are proline substitutions which likely destabilize the central N-SH2 β-sheet. (**Figure 6B**). Surprisingly, N-SH2 core mutants are rare in human diseases. Moreover, of the core mutants that can be accessed from wild-type SHP2 by a single nucleotide substitution, the few that are clinically-observed tend to be modestly activating (**Figure 6C**). For example, one of the few documented disease mutations in the N-SH2 core, L43F, only increases basal SHP2 activity three-fold (**Supplementary Table 4**). By contrast, the strongly activating L43P and L43R mutations have not been documented in clinical samples (**Figure 6C**).

**Figure 6.**
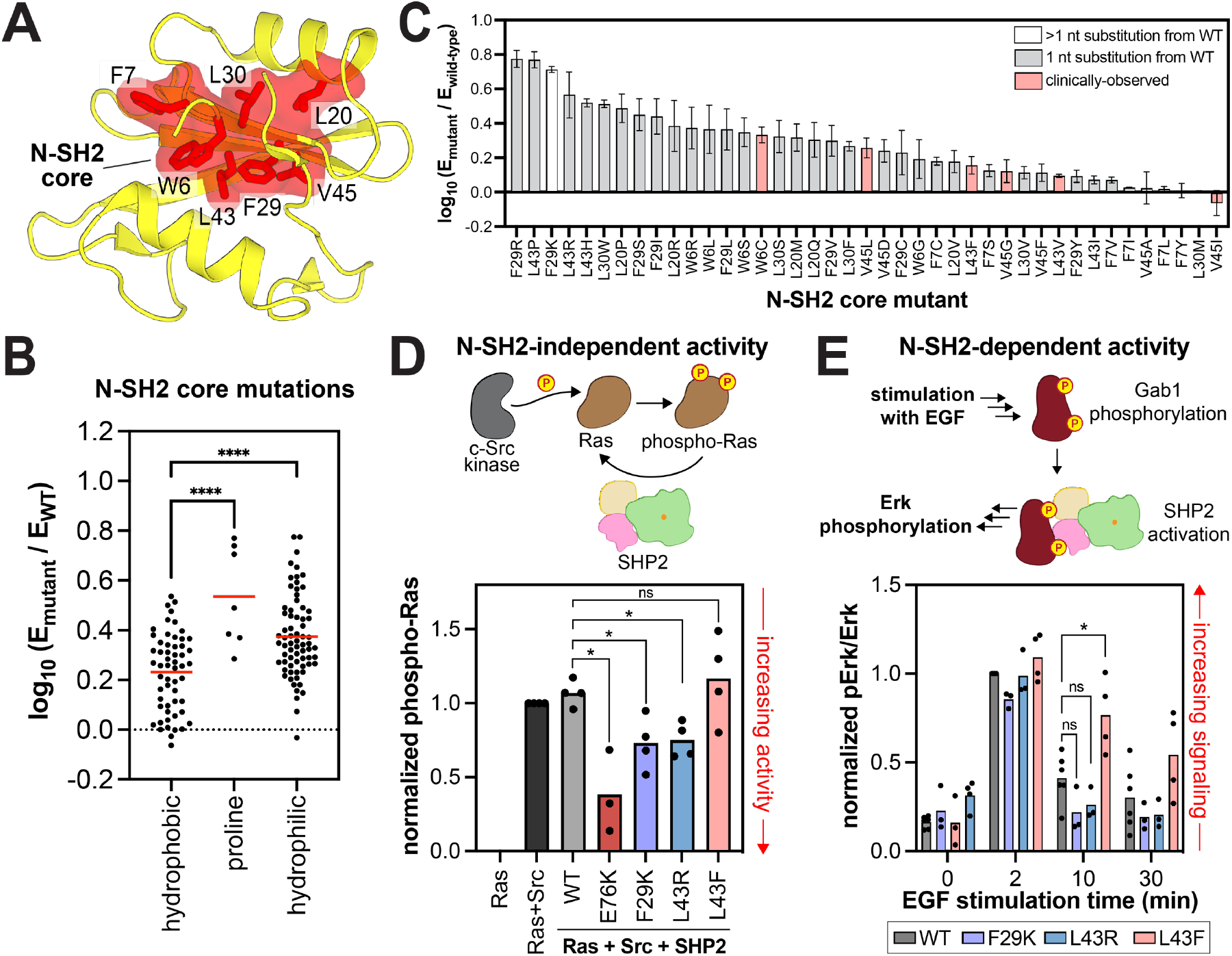
Functional effects and pathogenicity of mutations in the N-SH2 domain core. (**A**) Hydrophobic core residues in the N-SH2 domain (PDB code 1AYB). (**B**) Distribution of mutational effects at the residues shown in panel A in the SHP2_FL_ + v-Src_FL_ selection. Hydrophilic amino acids include C, D, E, H, K, N, Q, R, S, and T; hydrophobic amino acids include A, F, G, I, L, M, V, W, Y. The red line indicates the mean enrichment for all mutations in each group. (**C**) Functional effects of select N-SH2 core mutations in the SHP2_FL_ + v-Src_FL_ selection. Mutations observed in clinical databases (pink bars) are generally not the most activating, although many highly activating mutations are genetically accessible (1 nucleotide substitution away, gray bars). The SHP2_FL_ + v-Src_FL_ selection was used for panels B and C because it has greater dynamic range over activating mutations than the SHP2_FL_ + c-Src_KD_ selection. (**D**) N-SH2-independent catalytic activity of SHP2 N-SH2 core mutants and E76K in a Ras dephosphorylation assay in HEK293 cells (n = 3). Phospho-Ras levels are normalized to total Ras levels and to phospho-Ras in the Ras+Src sample. (**E**) Downstream signaling activity of SHP2 N-SH2 core mutants, dependent on N-SH2 binding to Gab1, in HEK293 cells stimulated with epidermal growth factor (EGF). Phospho-Erk levels are shown, normalized to total Erk levels and to the 2 minute time point for the wild-type SHP2 sample (n = 3-4). In panels B, D, and E, * denotes P < 0.05, **** denotes P < 0.0001.

We hypothesized that hyper-destabilizing N-SH2 core mutations such as L43R activate SHP2 at the expense of phosphoprotein binding capability, which may be expendable in our yeast selection assay but not in the context of pathogenic human signaling pathways. We were unable to purify SHP2_FL_ proteins or isolated N-SH2 domains containing polar substitutions in the hydrophobic core, likely because these mutations are severely destabilizing. Thus, to test our hypothesis, we expressed and assayed the hyperactive F29K and L43R mutants alongside the mildly-activating L43F mutant in two cellular assays for SHP2 activity, one that depends on N-SH2 function and one that does not. First, we examined the ability of the SHP2 mutants to dephosphorylate N-Ras that has been phosphorylated by co-expressed Src kinase^54^. This activity is not thought to depend on SH2 interactions (**Figure 6D**). Both F29K and L43R dephosphorylated N-Ras more efficiently than wild-type SHP2, whereas L43F was comparable to wild-type SHP2, consistent with the mutational scanning data (**Figure 6D and Supplementary Figure 6A**).

Next, we tested the signaling capabilities of these mutants in a context where N-SH2 binding functions are operative. SHP2 was co-expressed with Gab1, a known binding partner, cells were stimulated with the epidermal growth factor, and Erk phosphorylation was monitored as a downstream marker of SHP2-mediated activation of this pathway^22^. We previously showed that canonical hyperactivating mutations in SHP2 like E76K dramatically enhance Erk phosphorylation in this assay^22^. By contrast, the F29K and L43R mutants, which also activate SHP2, did not enhance Erk phosphorylation over wild-type SHP2, despite their increased catalytic activities (**Figure 6E and Supplementary Figure 6B**). The L43F disease mutant, on the other hand, modestly enhanced Erk phosphorylation. This mutant is destabilizing but can be purified (**Supplementary Figure 6C,D**), and it is still competent to bind phosphoproteins^22^. These results suggest that while the more disruptive N-SH2 core mutants display higher basal activity, their N-SH2 domains are too unstable to facilitate interactions with binding partners. Consequently, these mutants are unable to drive pathogenic signaling in human diseases. This balance between the disruption of SHP2 auto-inhibition and binding to upstream phosphoproteins constrains the spectrum of possible pathogenic mutations in the N-SH2 domain.

### SHP2 is dysregulated in human diseases through diverse structural perturbations

Our deep mutational scans allow us to classify SHP2 mutational effects into several distinct mechanisms of dysregulation, some of which were not previously known (**Figure 7**). Analysis of previously uncharacterized clinical variants in this framework can provide a more nuanced understanding of their potential pathogenicity. The classical mechanism of SHP2 dysregulation involves hyperactivation by mutations at N-SH2/PTP auto-inhibitory interface (*boxes 1 and 2*). On the PTP domain side of this interface, some mutations yield an open conformation at the expense of intrinsic phosphatase activity (*box 3*). Notably, several loss-of-activity mutants with disrupted auto-inhibition are frequently seen in diseases, suggesting a phosphatase-independent role of SHP2. Mutations can also dysregulate SHP2 function by altering the dynamics of the active-site WPD loop. Mutations directly on the WPD loop drive loop closure, dramatically enhancing intrinsic catalytic activity and simultaneously disrupting auto-inhibition (*box 4*). Other nearby mutations allosterically impact WPD loop dynamics to enhance intrinsic catalytic activity of the PTP domain, but their effects are not strong enough to disrupt auto-inhibition. These mutants most likely can still enhance signaling in the presence of the presence of activating phosphoprotein ligands (*box 5*). SHP2 can also be dysregulated by C-SH2/PTP interface mutations, which control the transition between SHP2 conformations (*box 6*). Finally, destabilization of the N-SH2 domain hydrophobic core can also disrupt auto-inhibition (*box 7*), but only those mutants with preserved phosphoprotein binding capabilities are likely to drive pathogenic signaling. We envision that this framework can be used to guide further mechanistic investigations into individual SHP2 variants.

**Figure 7.**
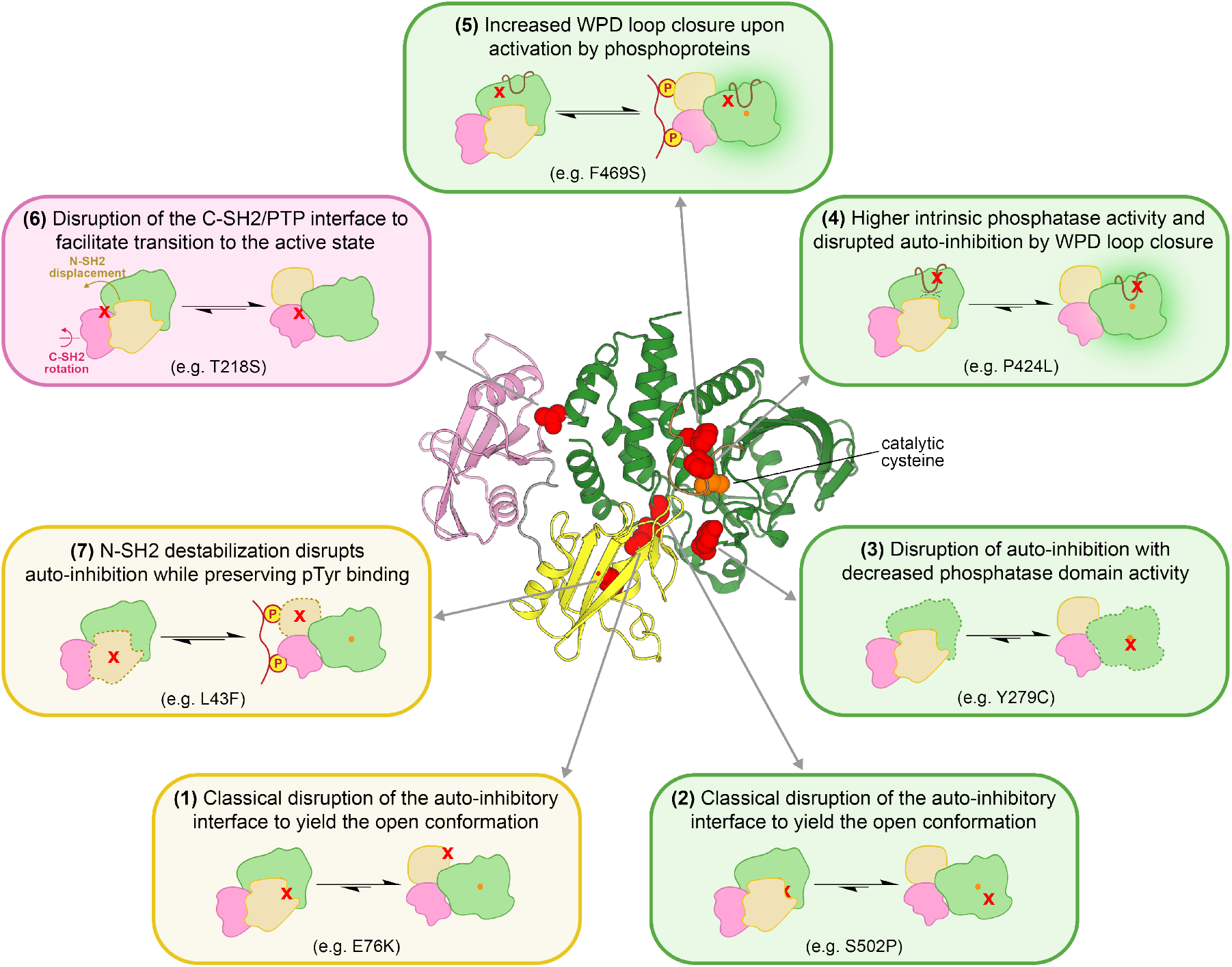
Diverse structural perturbations that can dysregulate SHP2 function. Representative mutations for each mechanism of dysregulation are highlighted on the auto-inhibited structure in red and specified in each box.

## Discussion

In this study, we used a yeast selection assay to compare the catalytic activities of nearly all possible point mutants in the human tyrosine phosphatase SHP2. We conducted deep mutational scans of both full-length SHP2 (593 residues, ∼12,000 variants) and its isolated catalytic domain (289 residues, ∼6000 variants). Thus far, our understanding of SHP2 regulation has largely been driven by the characterization of 10-20 disease-associated mutations, most of which cluster at the N-SH2/PTP interface^23,24,45^. For some of these well-studied disease mutants that are located on the phosphatase domain, comparative characterization of the full-length SHP2 constructs and the isolated PTP domains have allowed for rigorous dissection of the contributions of auto-inhibition and intrinsic phosphatase activity to overall enzyme dysregulation^24,42,45^. Our mutational scanning approach adds new depth to these analyses by enabling comparative mechanistic dissection across all PTP domain residues. More broadly, our datasets provide functional and mechanistic information on roughly 600 clinically observed SHP2 mutants, some within and some outside of the PTP domain, the vast majority of which were previously uncharacterized.

We focused our analysis on sets of mutations with significant but divergent activating or deactivating effects in either full-length SHP2 or the isolated PTP domain (**Figure 2G**). While our datasets do not directly assess pathogenicity, we conclude that mutants in each set dysregulate SHP2 via distinct mechanisms and are likely to affect SHP2-mediated signaling pathways. Further studies of the phenotypic effects of these mutations will provide more valuable insight into their potential pathogenicity. On the other hand, mutants with a neutral effect on basal activity, which our method primarily measures, are not necessarily benign. For example, we previously reported that the pathogenic N-SH2 mutant T42A alters phosphoprotein binding specificity, thereby sensitizing SHP2 to activation by specific proteins^22^. Moreover, our mutational scans captured the activating effects of pathogenic mutants that lack clear mechanistic explanations, such as the N-SH2/C-SH2 linker mutation D106A, and C-SH2 mutations R138Q and E139D (**Figure 2 and Supplementary Figure 2**). Alternative analytical methods will be required to dissect their pathogenicity.

Juxtaposition of our mutational scanning datasets with MD simulations allowed us to gain insights into the dynamics of two critical regions of SHP2: the C-SH2/PTP interface and the WPD loop. These analyses shed light on how these regions contribute to SHP2 regulation and pathogenicity. They also demonstrate how SHP2 diverges in its regulation from other PTP family members. For example, the role of T356 in governing WPD loop movement only exists in SHP2 and potentially PTPN9 (**Supplementary Figure 5D**), and the C-SH2/PTP interface SHP2, which governs activation and active state stability, diverges from that of its paralog SHP1 (**Supplementary Figure 7**)^55^. These variations in regulation and dynamics likely drive the broad range of catalytic activity and signaling functions seen across the PTP family. Finally, we note that our selection platform can easily be adapted to investigate mutational effects in other protein tyrosine phosphatases^37,38^. We envision that this platform, particularly when coupled with MD simulations, experimental biochemistry/biophysics, and cell signaling studies, will become invaluable tool to map sequence-structure-function relationships across this important enzyme family.

## Acknowledgements

We would like to thank the members of the Shah lab for their scientific insights and helpful discussions throughout the development of this project. We thank So Jung Lee from the Rothstein Lab for guidance with yeast work. We thank Jeanine Amacher for constructive feedback on this manuscript. This research was funded by NIH grant R35 GM138014 and American Cancer Society grant RSG-23-1038049-01-TBE to NHS. This work used the Expanse GPU cluster at the San Diego Supercomputer Center through allocation BIO220139 from the Advanced Cyberinfrastructure Coordination Ecosystem: Services & Support (ACCESS) program, which is supported by an NSF grants #2138259, #2138286, #2138307, #2137603, and #2138296.

