## Supplementary Information and Figures for "Deep mutational scanning of a multi-domain signaling protein reveals mechanisms of regulation and pathogenicity"

<u>**Supplementary Information for:**</u>


**Table of contents:**


Materials and Methods pg 2

Supp. Fig. 1. Yeast growth and selection by co-expression of Src kinase and SHP2 variants. pg 8

Supp. Fig. 2. Scanning mutagenesis and selection assays with $SHP2_{FL}$ and $SHP2_{PTP}$. pg 9

Supp. Fig. 3. Compensation for loss of phosphatase activity via disruption of auto-inhibition. pg 10

Supp. Fig. 4. Mutational sensitivity and dynamics at the C-SH2/PTP interface. pg 11

Supp. Fig. 5. PTP domain mutations that modulate WPD loop structure and dynamics. pg 12

Supp. Fig. 6. Destabilization of the N-SH2 domain by core mutations alters SHP activity. pg 13

Supp. Fig. 7. Divergent C-SH2/PTP interface interactions in SHP1 and SHP2. pg 14

Supplementary References pg 15

### Materials and Methods

#### Data availability

Deep sequencing data and molecular dynamics trajectory files will be made available via the following Dryad repository: https://doi.org/10.5061/dryad.83bk3jb18.

#### SHP2 mutagenesis library preparation

Saturation mutagenesis libraries of SHP2 were prepared with Mutagenesis by Integrated TilEs (MITE) method[1]. The 1782 bp (593AA + stop codon) full-length SHP2 gene we used was first optimized to yeast favorable codons. Then, the full-length sequence was divided into 15 separate tiles, each spanning around 40 amino acids (**Supplementary Table 1**). Two saturation mutagenesis oligo pools of alternating tiles were designed and acquired from Twist Bioscience. Each single amino acid substitution was encoded by a single oligo, thus there is no degeneracy in the library. The oligo sequences include invariant overhang sequences on each end designed for Gibson assembly to replace the wild-type sequence of each tile of a pET-28 plasmid containing the yeast-optimized human $SHP2_{FL}$ coding sequence. With PCR primers annealing to their overhang sequences, individual tiles were amplified from their oligo pool. Backbone amplification primers complementary to the tile amplification primers were used to amplify Gibson backbone DNA for each tile from the wild-type pET-28 yeast-optimized $SHP2_{FL}$ plasmid. Tile mutagenesis library inserts were then cloned onto their corresponding backbones using Gibson assembly to generate 15 separate plasmid libraries, each containing all single mutants of one tile region. For each tile, SHP2 full-length construct was PCR amplified with primers annealing right outside of the SHP2 gene on the plasmid, carrying overhangs for homologous recombination onto the yeast expression plasmid PWJ1781. For tiles 7-13 in the PTP domain, isolated PTP domain constructs spanning residues 235-539 were also amplified with homologous recombination overhangs.

#### Yeast expression plasmids construction and transformation

For expression of the kinase and phosphatase libraries in yeast, we used a galactose induced expression plasmid PWJ1781[2]. For the purposes of double transformation in yeast, we switched the LEU2 marker to a URA3 marker to generate PWJ1781-URA. v-$Src_{FL}$ and c-$Src_{KD}$ DNA sequences were then integrated into PWJ1781-URA by Gibson assembly to make kinase expression plasmids PWJ-1781-URA-c-$Src_{KD}$ and PWJ-1781-URAv-$Src_{FL}$. SHP2 expression plasmids were constructed with homologous recombination in yeast for both the $SHP2_{FL}$ construct and $SHP2_{PTP}$. In both cases, inserts bearing each mutagenized tile were integrated into PWJ1781 separately. PWJ1781 was first digested with Hpa1 to yield linear backbone DNA. Then, a molar ratio 1:2 of digested backbone and amplified insert was co-transformed into yeast strain YPH499 with the LiAc/PEG/ssDNA method[3]. Specifically, 2 µg of the two DNA pieces, total, were transformed into ~3 x $10^8$ YPH499 cells. The transformed cells were then grown in 500 mL synthetic complete media without leucine supplemented with 4% glucose at 30 °C with shaking for ~40 hours to reach a high concentration ($OD_{600}$ ~6-10, 6-10 x $10^7$ cells/mL). The cells from each transformation were collected as stock for one tile. Cells from each stock were directly subjected to kinase transformation. In the kinase plasmid transformation, 2 µg of PWJ-1781-URA-c-$Src_{KD}$ and PWJ-1781-URAv-$Src_{FL}$ was transformed into ~6 x $10^7$ YPH499 cell bearing phosphatase plasmid with the LiAc/PEG/ssDNA method. The doubly transformed cells were grown in 100 mL of synthetic complete media without leucine and uracil supplemented with 4% glucose at 30 °C with shaking for ~28 hours to reach a high OD ($OD_{600}$ ~4, 4 x $10^7$ cells/mL). Each transformation yielded a cell stock transformed with one of the v-$Src_{FL}$ or c-$Src_{KD}$ expression plasmids and one of the $SHP2_{FL}$ or $SHP2_{PTP}$ plasmids bearing one mutagenized tile. Each stock was then subjected to outgrowth and selection individually.

#### Tile library selection with the yeast growth assay

Each cell stock bearing one kinase expression plasmid and a SHP2 library with one variable tile was inoculated into synthetic complete media without uracil and leucine supplemented with glycerol lactate (2% lactic acid, 3% glycerol, 0.05% glucose in media) at a starting $OD_{600}$ = 0.1. The culture was grown at 30 °C with shaking for ~16 hours and then used to inoculate synthetic complete media without uracil and leucine supplemented with 4% galactose for co-expression and selection. The rest of the cells grown in glycerol lactate media were harvested as unselected samples. For each outgrowth culture, two parallel selection cultures were made with a starting $OD_{600}$ = 0.05. The cultures were allowed to grow in 30 °C with shaking for 24 hours, and the cells were then harvested as post-select samples.

#### Deep sequencing

For cell stocks with each tile before and after selection, plasmid DNA was extracted with Zymoprep Yeast Plasmid Miniprep II kit from ~1 x $10^8$ cells. Each mutagenized tile DNA library was PCR amplified from its corresponding SHP2 plasmid with tile specific primers bearing overhangs for the addition of Illumina sequencing adapters. The PCR mix was directly followed by another round of PCR amplification with Illumina barcoding primers to append Illumina sequencing adaptors and 5' and 3' indices (D700 and D500 series primers). The PCR products were gel purified and quantified with QuantiFluor® dsDNA System (Promega). Then, samples were pooled at ratios that all mutants are theoretically equally represented, and the pooled libraries were sequenced on a MiSeq using V2 300 cycle reagent kits. Each sequencing run contained no more than 24 samples to ensure good read counts.

#### Sequencing data analysis

Paired-end sequencing reads were first merged with FLASH[4], followed by trimming with Cutadapt to remove constant sequences outside of the mutagenized libraries[5]. Then, read counts for each mutant in the libraries were calculated using in-house Python scripts (https://github.com/nshahlab/2024_Jiang-et-al_SHP2-DMS). For each sequenced library, frequencies of the mutants ($f_{mut}$) were first calculated by taking the ratio of the mutants' read counts ($n_{mut}$) over total reads ($n_{total}$) in the library (euation 1). Then, enrichment ($E_{mut}$) were calculated through dividing after selection ($f_{selected}$) by frequencies before selection ($f_{unselected}$) (equation 2). The enrichment scores data that we are presenting on the heat maps are $log_{10}$-transformed enrichment normalized to WT (equation 3). The distribution of enrichment scores from selection in the $SHP2_{FL}$ context was fit to a Gaussian distribution, and datapoints outside of $\mu \pm 2\sigma$ were defined as significantly gain- or loss-of-activity. The distribution of enrichment scores from selections in the $SHP2_{PTP}$ context followed a bimodal distribution and was fit to the sum of two Gaussians. Datapoints outside of $\mu \pm 2\sigma$ for the distribution around 0 were defined as significantly gain- or loss-of-activity.

(1) $f_{mut} = \frac{n_{mut}}{n_{total}}$ (2) $E_{mut} = \frac{f_{selected}}{f_{unselected}}$ (3) $Score_{mut} = \log_{10} E_{mut} - \log_{10} E_{WT}$

#### Purification of full-length SHP2 proteins

pET28-His-TEV plasmid encoding the human $SHP2_{FL}$ sequence (human cDNA) was used for QuikChange mutagenesis to generate $SHP2_{FL}$ mutants.[6] For purification of wild-type and mutant $SHP2_{FL}$ constructs, plasmids were first transformed into chemically competent BL21(DE3) cells and grown on LB agar plates supplemented with 50 µg/mL kanamycin. Then, colonies scraped off the plates were inoculated into 100 mL LB with kanamycin and grew at 37 °C to $OD_{600}$ = 1. 50mL of this culture was used to inoculate 1 L LB cultures with kanamycin at a starting $OD_{600}$ = 0.1, and the 1 L cultures were incubated at 37 °C until their $OD_{600}$ reached 0.5. 0.5 mM IPTG was supplemented to induce expression, and the expression cultures were grown overnight at 18 °C. The cultures were then spun down at 4000xg for 30min, and resuspended in lysis buffer (50 mM Tris pH 8.0, 300 mM NaCl, 20 mM imidazole, 10% glycerol, and freshly added 2 mM β-mercaptoethanol). The cell suspensions were lysed using sonication (Fisherbrand Sonic Dismembrator), and spun down at 14,000 x g for 45 minutes. The His-tagged $SHP2_{FL}$ constructs were extracted from the supernatant with a 5 mL Ni-NTA column (Cytiva). The column was subsequently washed with 50 mL lysis buffer and 50 mL wash buffer (50 mM Tris pH 8.5, 50 mM NaCl, 20 mM imidazole, 10% glycerol, and freshly added 2 mM β-mercaptoethanol), followed by elution of the tagged SHP2 with a mixture of 25 mL wash buffer + 25 mL elution buffer (50 mM Tris pH 8.5, 50 mM NaCl, 500 mM imidazole, 10% glycerol) directly onto a 5 mL HiTrap Q Anion exchange column (Cytiva). The Q column was washed once with 40 mL anion exchange buffer A (50 mM Tris pH 8.5, 50 mM NaCl, 1 mM TCEP), and the protein was eluted off with a salt gradient between Anion A buffer and Anion B buffer (50 mM Tris pH 8.5, 1 M NaCl, 1 mM TCEP). The eluted protein fractions were collected and cleaved with 0.10 mg/mL $His_6$-tagged TEV protease at 4 °C overnight to remove the His tag. The cleavage mixture was applied through 2 mL of Ni-NTA gravity column (ThermoFisher) to remove uncleaved protein and TEV protease, and the flow through was concentrated to less than 1 mL. Finally, the concentrated protein solution was loaded onto a Superdex 200 16/600 gel filtration column (Cytiva) equilibrated with SEC buffer (20 mM HEPES pH 7.5, 150 mM NaCl, and 10% glycerol) for size exclusion purification. Pure fractions were pooled, concentrated, and flash frozen in liquid $N_2$ for storage at −80 °C.

#### Purification of SHP2 SH2 domains

pET28-His6-SUMO-NSH2-Avi and pET28-His6-SUMO-CSH2-Avi plasmids in our lab were used as templates for wild-type and mutant SH2 domain purification[6]. The Avi tags were first removed in one cloning step, and the resulting pET28-His6-SUMO-N/C-SH2 plasmids were applied to QuikChange mutagenesis to generate desired SH2 mutants. For purification of wild-type and mutant SH2 constructs, plasmids were first transformed into chemically competent BL21(DE3) cells and grown on LB agar plates supplemented with 50 µg/mL kanamycin. Then, colonies scraped off the plates were inoculated into 100 mL LB with kanamycin and grew at 37 °C to $OD_{600}$ = 1. 50 mL of this culture was used to inoculate 1 L LB cultures with kanamycin at a starting $OD_{600}$ = 0.1, and the 1L cultures were at 37 °C until their OD reached 0.5. 0.5mM IPTG was supplemented to induce expression, and the expression cultures were grown overnight at 18 °C. The cultures were then spun down at 4000 x g for 30 min, and resuspended in lysis buffer (50 mM Tris pH 7.5, 300 mM NaCl, 20 mM imidazole, 10% glycerol, and freshly added 2 mM β-mercaptoethanol). The cell suspensions were lysed using sonication (Fisherbrand Sonic Dismembrator), and spun down at 14,000xg for 45 minutes. The His-tagged SH2 constructs were extracted from the supernatant with a 5 mL Ni-NTA column (Cytiva). The column was subsequently washed with 50 mL lysis buffer and 50 mL wash buffer (50 mM Tris pH 7.5, 50 mM NaCl, 20 mM imidazole, 10% glycerol, and freshly added 2 mM β-mercaptoethanol), followed by elution of the tagged SH2 with a mixture of 25mL wash buffer + 25mL elution buffer (50 mM Tris pH 7.5, 50 mM NaCl, 500 mM imidazole, 10% glycerol) directly onto a 5 mL HiTrap Q Anion exchange column (Cytiva). The Q column was washed once with 40 mL anion exchange buffer A (50 mM Tris pH 7.5, 50 mM NaCl, 1 mM TCEP), and the protein was eluted off with a salt gradient between Anion A buffer and Anion B buffer (50 mM Tris pH 7.5, 1 M NaCl, 1 mM TCEP). The eluted protein fractions were collected and cleaved with 0.05mg/mL $His_6$-tagged Ulp1 protease at 4°C overnight to remove the His tag. The cleavage mixture was applied through 2mL of Ni-NTA gravity column (ThermoFisher) to remove uncleaved protein and TEV protease, and the flow through was concentrated to less than 1mL. Finally, the concentrated protein solution was loaded onto a Superdex 75 16/600 gel filtration column (Cytiva) equilibrated with SH2-SEC buffer (20 mM HEPES pH 7.4, 150 mM NaCl, and 10% glycerol) for size exclusion purification. Pure fractions were pooled and concentrated, and flash frozen in liquid $N_2$ for long-term storage at −80°C.

#### SHP2 basal activity measurements

Basal activities of wild-type and mutant SHP2 were measured against the fluorogenic substrate 6,8-difluoro-4-methylumbelliferyl phosphate (DiFMUP). Initial DiFMUP dephosphorylation rates by SHP2 variants were measured at 37 °C in triplicate. Reactions were done in black polystyrene flat bottom half area 96-well plates at a working volume of 50 µL. For each replicate, the reaction mix contains a $SHP2_{FL}$ concentration of 0.5-3.0 nM, depending on the variant, and a DiFMUP concentration series of 4000, 2000, 1000, 500, 250, 125, 62.5 and 31.25 µM. With each plate the absorbance of the dephosphorylation product DiFMU at a concentration series of 200, 100, 50, 25, 12.5, 6.25, 3.125 and 0 µM was measured as a standard curve to convert absorbance values to product concentrations. Reactions were started by the addition of the protein, and emitted fluorescence at 455 nm was measured every 25 seconds within 50 min with a BioTek Synergy Neo2 multi-mode reader. Fluorescence values were converted into DiFMU concentrations, and initial rates were determined by the slope of the first 5 minutes on the reaction curves. All the initial rates were fitted onto Michaelis-Menten curves using GraphPad Prism to determine $k_{cat}$ and $K_M$ values.[7] Catalytic Parameters are compiled in **Supplementary Table 4**.

#### Melting temperature measurements via differential scanning fluorimetry (DSF)

Melting temperature measurements were conducted in DSF buffer (20 mM HEPES pH 7.5, 50 mM NaCl, 0.4% DMSO) on MicroAmp Fast Optical 96-well Reaction plates (Applied Biosystems, # 4346906) at working volumes of 20µL. The mixtures contained 10 µM protein and 25x SYPRO Orange Protein Gel Stain (Thermo Fisher, catalog no. S-6650). Melting curves were measured on an Applied Biosystems Step-One Plus RT-PCR thermocycler between 25 °C and 95 °C with a gradient of +0.5 °C per minute (excitation: 472 nm; emission: 570 nm). Fluorescence reads under each temperature were analyzed using DSFworld and melting temperatures were calculated with dRFU[8].

#### General cell culture

HEK 293 cells were grown at 37 °C with 5% $CO_2$ in Dubecco's Modified Eagle Medium (DMEM) supplemented with 10% Fetal Bovine Serum (FBS).

#### EGF stimulation experiments

2.2 x $10^6$ HEK 293 cells were seeded in a 10 cm plate. The next day, cells were transfected overnight with 5 µg Gab1 and 5µg SHP2 using 30 µg of polyethyleneimine in 1 mL DMEM. Transfection media was refreshed and cells were serum-starved for 24 hours in DMEM. Cells were harvested by scraping, washed 3 times in 1 mL PBS at room temperature. Prior to stimulation, an aliquot of cells was taken as an unstimulated control (t = 0). Cells were then resuspended in 25 ng/mL EGF in pre-heated PBS and placed in a 37 °C heatblock. Aliquots were taken at 2, 10 and 30 minutes; placed on ice and spun down in a 4 °C tabletop centrifuge at 1000 g for 5 minutes. Supernatant was aspirated and cells were lysed in 75 µL lysis buffer (20 mM Tris pH 8.0, 137 mM NaCl, 2 mM EDTA, 10% glycerol, 0.5% NP-40, with freshly added phosphatase- and protease inhibitors) for 25 minutes on ice. Lysates were spun down for 15 minutes at 17,000 g in a 4 °C tabletop centrifuge. Supernatant was used in a bicinchoninic acid (BCA) assay to determine protein concentration. 15 µg of total protein was loaded on a 12% acrylamide gel and transferred to a 0.45 µm nitrocellulose membrane using the StandardSD protocol on the Bio-Rad Trans-Blot Turbo. Membranes were blocked for 1 hour at room temperature using 5% Bovine Serum Albumin (BSA) in TBS. Primary antibodies were stained overnight at 4 °C (Erk 1:1000, p-Erk 1:2000, Vinculin 1:1000, Myc 1:5000, FLAG 1:5000) in 5% BSA in TBST. Membranes were washed 3 times in 5 mL TBST for 5 minutes each. Secondary antibodies were incubated in 5% BSA in TBST for 1 hour at room temperature (1:10,000). Membranes were imaged on a LiCor Odyssey and bands were quantified using ImageStudio.

#### Ras dephosphorylation assay (Rassay)

0.8 x $10^6$ HEK 293 cells were seeded in a 6 cm plate. The next day, cells were transfected with Ras, Ras and Src, or Ras, Src and SHP2; to a total of 3 µg (a pEF vector with no open reading frame was used to make up the difference between conditions) in 300 µL DMEM with 9 µg of polyethyleneimine. Transfection medium was refreshed the next morning and replaced with warm DMEM with 10% FBS. 36 hours after transfection, cells were harvested by scraping and washed 3 times in 1 mL cold PBS. Cells were lysed in 150 µL lysis buffer (20 mM Tris pH 8.0, 137 mM NaCl, 2 mM EDTA, 10% glycerol, 0.5% NP-40, with freshly added phosphatase- and protease inhibitors) for 25 minutes on ice. Lysates were spun down for 15 minutes at 17,000 g in a 4 °C tabletop centrifuge. Supernatant was used in a BCA assay to determine protein concentration. 80 µg of protein was used in an immuno-precipitation (IP) with 5 µg of packed Pierce anti-HA magnetic beads (Fisher, #88836) in a total volume of 350 µL. Samples were incubated at 4 °C overnight while rotating. The next morning, beads were washed 3 times on a magnetic racks using 1 mL of lysis buffer, and finally resuspended in 65 µL of 1x Laemmli buffer. All samples were boiled at 100 °C for 8 minutes, and 15 µg of total protein (total cell lysate) or 15 µL of each sample (IP) was loaded onto a 12% acrylamide gel. Proteins were transferred to a 0.45 µm nitrocellulose membrane using the StandardSD protocol on the Bio-Rad Trans-Blot Turbo. Membranes were blocked for 1 hour at room temperature using 5% BSA in TBS. Primary antibodies were stained for 2 hours at room temperature (TCL: Src 1:1000, β-actin 1:5000, Myc 1:5000, HA 1:1000; IP: HA 1:1000, pTyr 1:2000) in 5% BSA in TBST. Membranes were washed 3 times in 5 mL TBST for 5 minutes each. Secondary antibodies were incubated in 5% BSA in TBST for 1 hour at room temperature (1:10,000). Membranes were imaged on a LiCor Odyssey and bands were quantified using ImageStudio.

The specific antibodies used for this study are listed in the table below:

| Description | Vendor | Catalog # |
|---|---|---|
| Vinculin | CST | 13901S |
| FLAG (Clone 2B3C4) | Fisher | 50-173-6451 |
| Myc (Clone (9E10) | Invitrogen | R95025 |
| p44/42 (Erk1/2) (L34F12) | CST | 4696S |
| p-p44/42 (p-Erk1/2) (197G2) | CST | 13214S |
| c-Src | CST | 2123S |
| B-actin | Sigma | A5441 |
| HA (TCL) | Sigma | SAB5600116 |
| Phospho-Tyrosine (P-Tyr-1000) MultiMab | CST | 8954S |
| HA (IP) | Sigma | 11867423001 |
| IRDye® 680RD Goat anti-Rabbit IgG | LiCor | 926-68071 |
| IRDye® 800CW Goat anti-Mouse IgG | LiCor | 926-32210 |
| IRDye 800CW Goat anti-Rat IgG | LiCor | 926-32219 |

#### Analysis of COSMIC, TCGA, and ClinVar databases

We extracted clinically observed SHP2 mutant data from COSMIC[9], TCGA[10], and ClinVar[11] (**Supplementary Table 5**). COSMIC and ClinVar data were downloaded in September 2023. TCGA data was downloaded in October 2024. All disease mutations extracted from the three databases combined are annotated as "clinically observed" in our analysis. Cancer mutations in TCGA and COSMIC are classified as "all cancer mutations", and cancer mutations with more than five reports are classified as "high frequency cancer mutations". Pathogenic mutations and variants of uncertain significance annotated in ClinVar are also separately classified, and enrichment scores of mutations in all categories were plotted to visualize their activity distributions (**Figure 3B,E**). Mutants in COSMIC are also classified based on primary tissue type, and enrichment scores of variants in each cancer type were plotted as well (**Figure 3C,F**)

#### Molecular dynamics simulations

##### *Preparation of Structural Models for Simulations*

We built and simulated near full-length SHP2 (residues 1-528) in the closed, auto-inhibited state, and in two active conformational states. For all three states, we considered the wild-type protein, as well as the protein with Glu 76 mutated to Lys. For the simulations of the protein in the auto-inhibited state, we used the crystal structure 4DGP as the starting structure[12]. Missing residues were modeled as follows. Residues 1-3 at the N-terminal end were built using PyMOL[13]. Residues 235-245 were taken from a model of the full-length SHP2 that was predicted by AlphaFold2[14], since this region is missing in all crystal structures. Residues 293-303 were taken from the crystal structure 6CRF[15] and residues 314-232 were taken from the crystal structure 4RDD[16]. For simulations of the protein with the E76K mutation, Glu 76 was mutated to Lys using PyMOL. Crystalline waters were retained in the starting structure. For the protein in the first open, active state, we used the crystal structure 6CRF as the starting structure[15]. In this structure Glu 76 is mutated to Lys, so for the wild-type simulations, this residue was mutated to Glu using PyMOL. Missing residues 89-93, 140-145, 154-166 and 203-209 were taken from 4DGP[12]. Residues 237-244 were taken from a model generated by AlphaFold2[14]. Residues 313-324 were taken from 4RDD[16], and missing C-terminal residues 526-528 were built in using PyMOL. The alternative active state model was generated by AlphaFold2, by inputting the wild-type SHP2 sequence (residues 1-528) into ColabFold with the default settings[17]. The N-SH2 domains are displaced in the AlphaFold2 structure when compared to the crystal structure (PDB code 6CRF), but they are otherwise similar. Crystalline waters are present in 6CRF. These waters were retained in the simulations starting from the crystal structure and were added to the AlphaFold2 structures. Cys 459 is deprotonated in all systems. Both N- and C-termini were capped with acetyl and amide groups, respectively, in all systems.

*Simulation Protocol*

Each system was solvated with TIP3P water[18], and ions were added such that the final ionic strength of the system was 100 mM using the tleap package in AmberTools22[19]. The energy of each system was minimized first for 5000 steps while holding the protein atoms and crystalline waters fixed, followed by minimization for 5000 steps while allowing all the atoms to move. Following minimization, three individual trajectories were generated for each system, with distinct initial velocities for each. The temperature of each system was raised in two stages – first to 100 K over 0.5 ns and then to 300 K over 0.5 ns. The protein atoms and crystalline waters were held fixed during the heating stage. Each system was then equilibrated for 2 ns, followed by production runs. Three production trajectories, each 2.5 μs long, were generated for each system. All equilibration runs and production runs were performed at constant temperature (300 K) and pressure (1 bar). The simulations were carried out with the Amber package[20] using the ff14SB force field for proteins[21] using an integration timestep of 2 fs. The Particle Mesh Ewald approximation was used to calculate long-range electrostatic energies[22]. All hydrogens bonded to heavy atoms were constrained with the SHAKE algorithm[23]. The Langevin thermostat was used to control the temperature with a collision frequency of 1 $ps^{-1}$. Pressure was controlled while maintaining periodic boundary conditions.

*Analyses*

MD trajectories were compiled from the raw data using the CPPTRAJ module of AmberTools22[19]. Structures were extracted from the trajectories in both 1 ns and 10 ns increments for analysis and visualization. All measurements and calculations were done using the PDB module in Biopython[24]. For most calculations reported in the main text, trajectories were sampled every 10 ns. The measurements from all three replicates of each system were combined to determine the reported distributions. In cases where distance calculations involved a redundant atom (e.g. distances between two possible nitrogens and two possible oxygens in a Glu/Arg ion pair), all combinations of distance measurements were calculated, then the shortest distance at each frame was determined and used for the distribution plots. For visualization, trajectories were sampled every 10 ns. All structure visualization and rendering in this study was done using PyMOL[13].

*Acknowledgements for computational resources*

This work used the Expanse GPU cluster at the San Diego Supercomputer Center through allocation BIO220139 from the Advanced Cyberinfrastructure Coordination Ecosystem: Services & Support (ACCESS) program, which is supported by an NSF grants #2138259, #2138286, #2138307, #2137603, and #2138296.

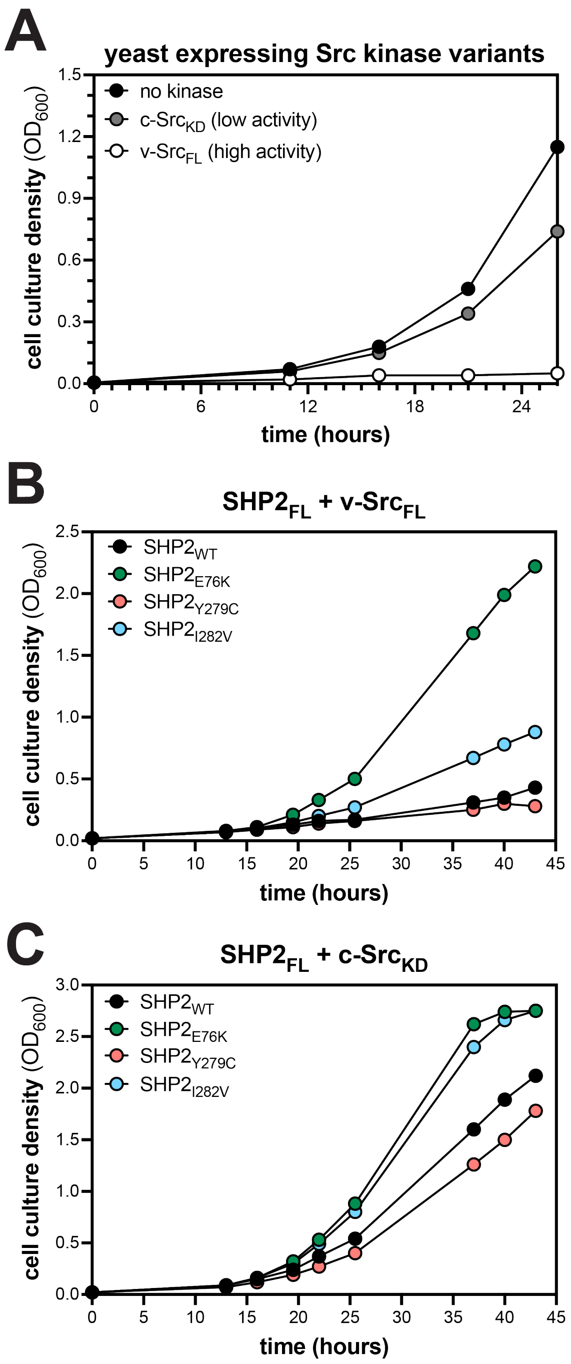


**Supplementary Figure 1. Yeast growth and selection by co-expression of Src kinase and SHP2 variants.** (**A**) Suppression of yeast growth by expression of v-Src$_{FL}$ and c-Src$_{KD}$. SHP2 was not expressed in these experiments. (**B**) Rescue of yeast growth from v-Src$_{FL}$ toxicity by co-expression of SHP2 variants. (**C**) Rescue of yeast growth from c-Src$_{KD}$ toxicity by co-expression of SHP2 variants.

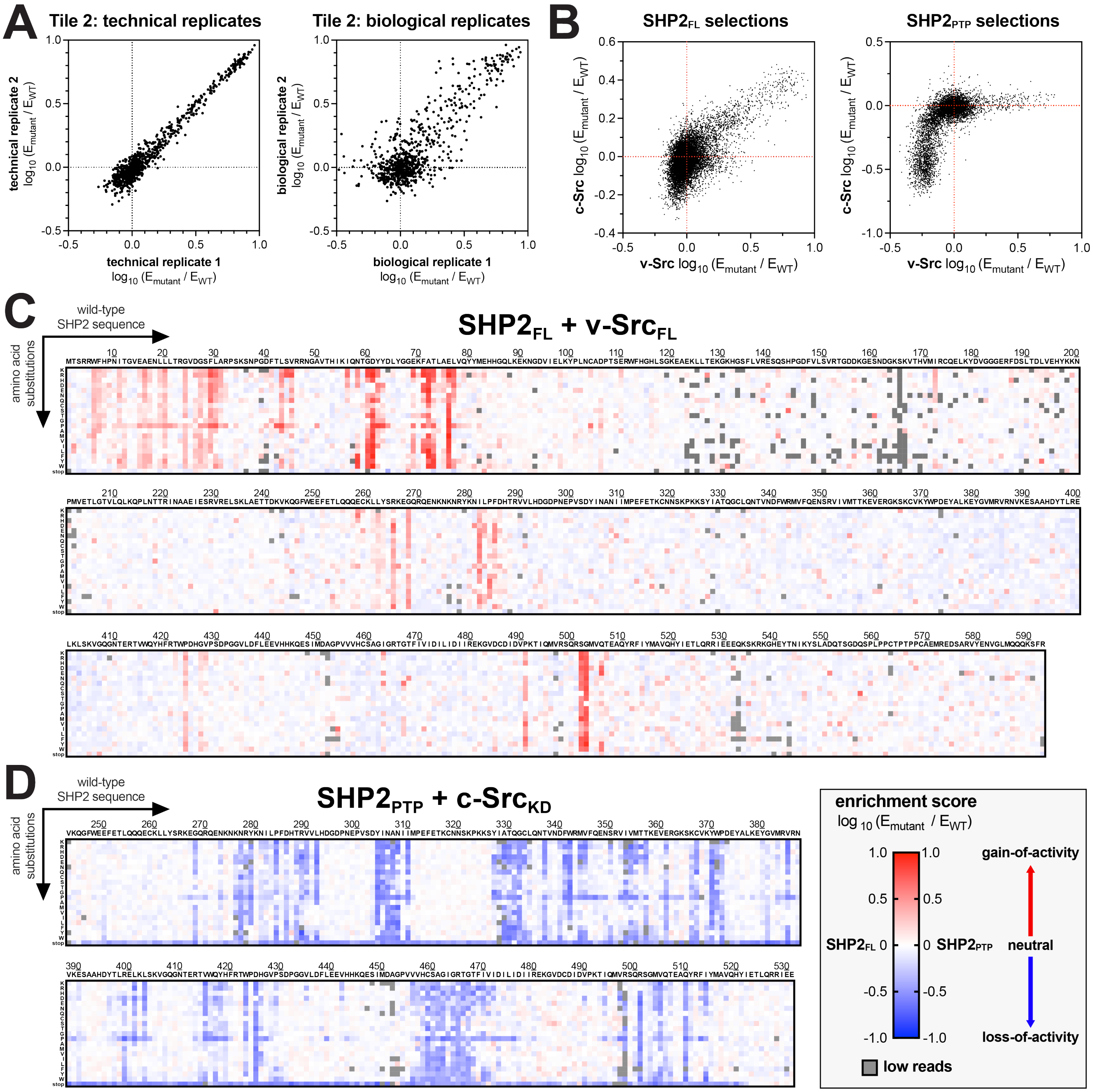


**Supplementary Figure 2. Scanning mutagenesis and selection assays with SHP2$_{FL}$ and SHP2$_{PTP}$.** (**A**) Representative replicate correlations for individual tiles. Tile 2 is used as an example. Comparisons are shown for two outgrowth and selection experiment done from a single transformation (technical replicates) and two outgrowth and selection experiments done from two different transformations (biological replicates). (**B**) Correlation between enrichment scores from the two SHP2$_{FL}$ mutational scans (*left*) and the two SHP2$_{PTP}$ mutational scans (*right*). (**C**) Heatmap depicting the enrichment scores for SHP2$_{FL}$ co-expressed with v-Src$_{FL}$ (n = 2-4). (**D**) Heatmap depicting the enrichment scores for SHP2$_{PTP}$ co-expressed with c-Src$_{KD}$ (n = 2). SHP2$_{FL}$ mutational scanning datasets, including clinical annotations, can be found in **Supplementary Table 2**. SHP2$_{PTP}$ mutational scanning datasets can be found in **Supplementary Table 3**.

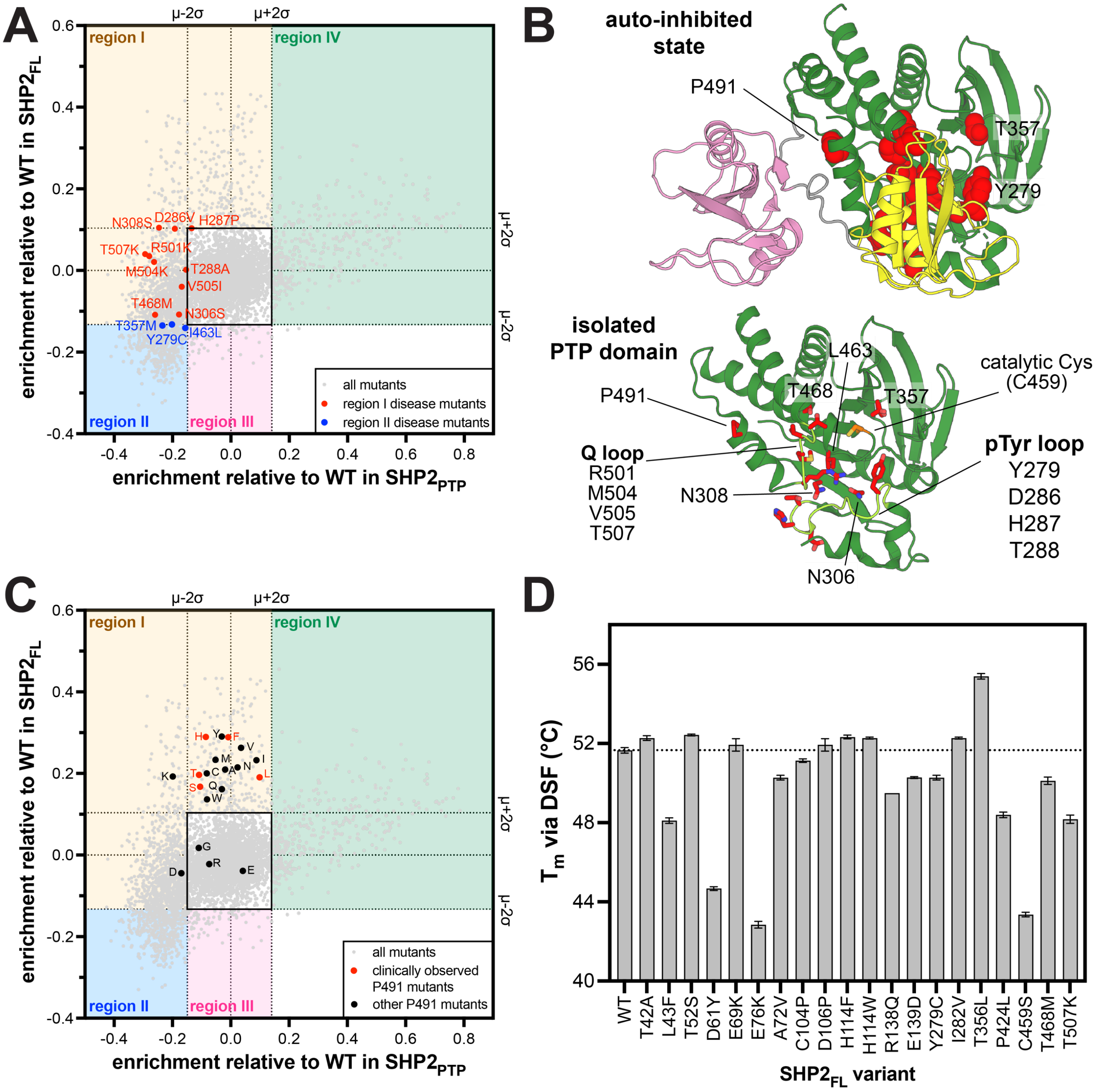


**Supplementary Figure 3. Compensation for loss of phosphatase activity by disruption of auto-inhibition.** (**A**) Scatterplot juxtaposing $\text{SHP2}_{FL}$ and $\text{SHP2}_{PTP}$ mutational scanning datasets, highlighting mutations of interest in regions I and II. Highlighted mutations in region I reduce intrinsic phosphatase activity but are near-neutral in a $\text{SHP2}_{FL}$ context, due to disrupted auto-inhibition. Highlighted mutations in region II reduce intrinsic phosphatase activity to a point that is beyond rescue by disruption of auto-inhibition. (**B**) Mutations in region I and II that disrupt auto-inhibition and in some cases also disrupt intrinsic phosphatase activity, highlighted on the auto-inhibited SHP2 structure (PDB code 4DGP) or a structure of the isolated PTP domain (PDB code 3ZM0). Most mutations are located at the N-SH2/PTP interface or on key active site loops. P491 is located at the C-SH2/PTP interface. (**C**) Scatterplot juxtaposing $\text{SHP2}_{FL}$ and $\text{SHP2}_{PTP}$ mutational scanning datasets, highlighting the high density of P491 mutations in and around region I. (**D**) Melting temperatures ($T_m$) for $\text{SHP2}_{FL}$ mutants, measured by differential scanning fluorimetry (DSF). $T_m$ is generally reduced for mutants that are less auto-inhibited.

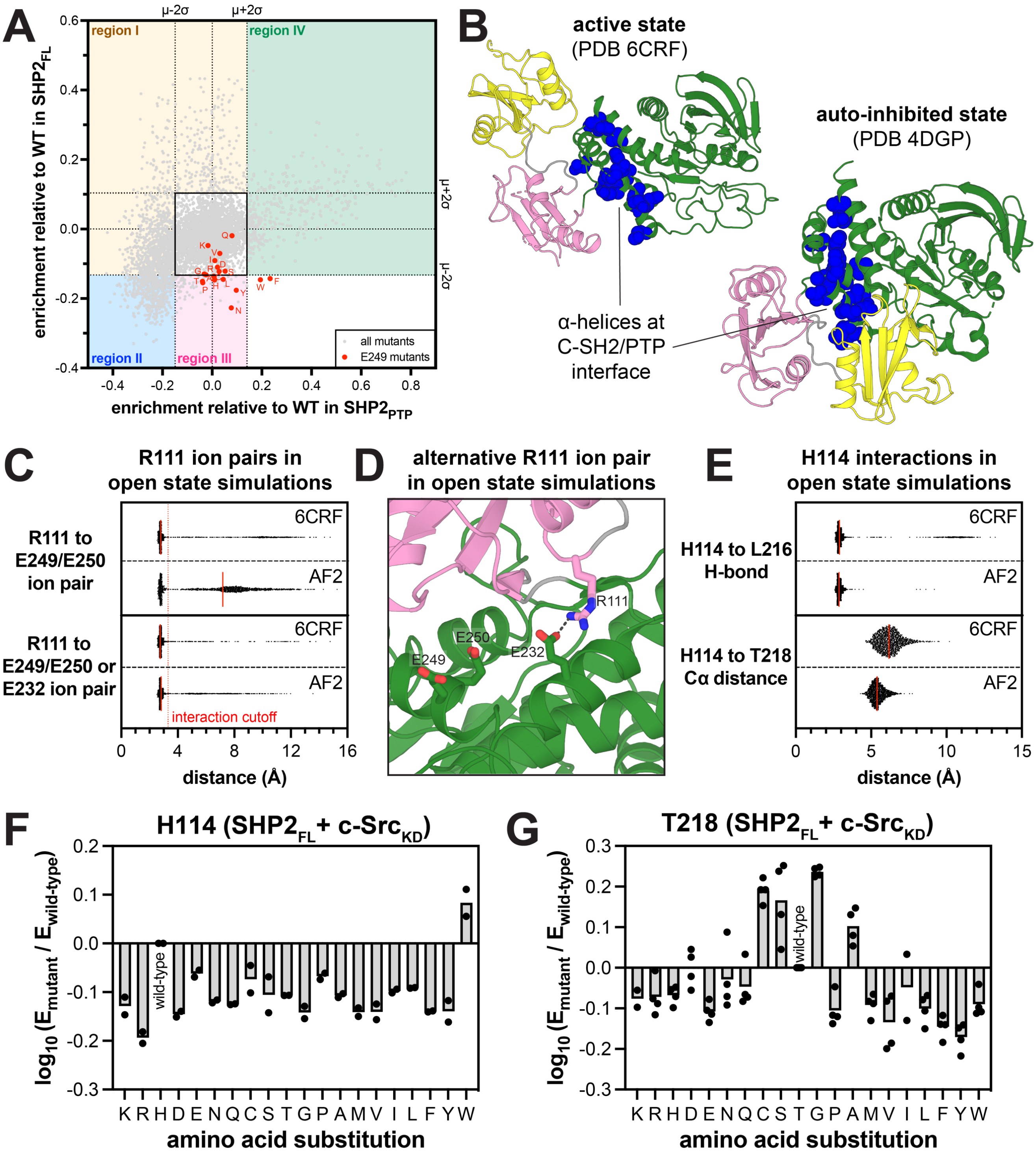


**Supplementary Figure 4. Mutational sensitivity and dynamics at the C-SH2/PTP interface.** (**A**) Scatterplot juxtaposing SHP2$_{FL}$ and SHP2$_{PTP}$ mutational scanning datasets, highlighting the high density of E249 mutations in and around region III. (**B**) Major deactivating mutation sites in region III at the C-SH2/PTP interface. (**C**) Distribution of distances in open state simulations for ion pairs with R111. Shortest distance to one of the primary pairing partners (E249 or E250) is plotted, as well as the shortest distance to E249, E250, or the secondary partner E232. Each distribution shows data for 6 simulations starting from 6CRF or the AlphaFold2 model. (**D**) Structure of R111 interacting with E232 in open simulations. (**E**) Distribution of distances in open state simulations for interactions between H114 and L216 (side-chain to backbone H-bond) and H114 and T218 (Cα-Cα). Each distribution shows data for 6 simulations starting from 6CRF or the AlphaFold2 model. (**F**) Mutational effects at H114 in the SHP2$_{FL}$ selection assays with c-Src$_{KD}$. (**G**) Mutational effects at T218 in the SHP2$_{FL}$ selection assays with c-Src$_{KD}$.

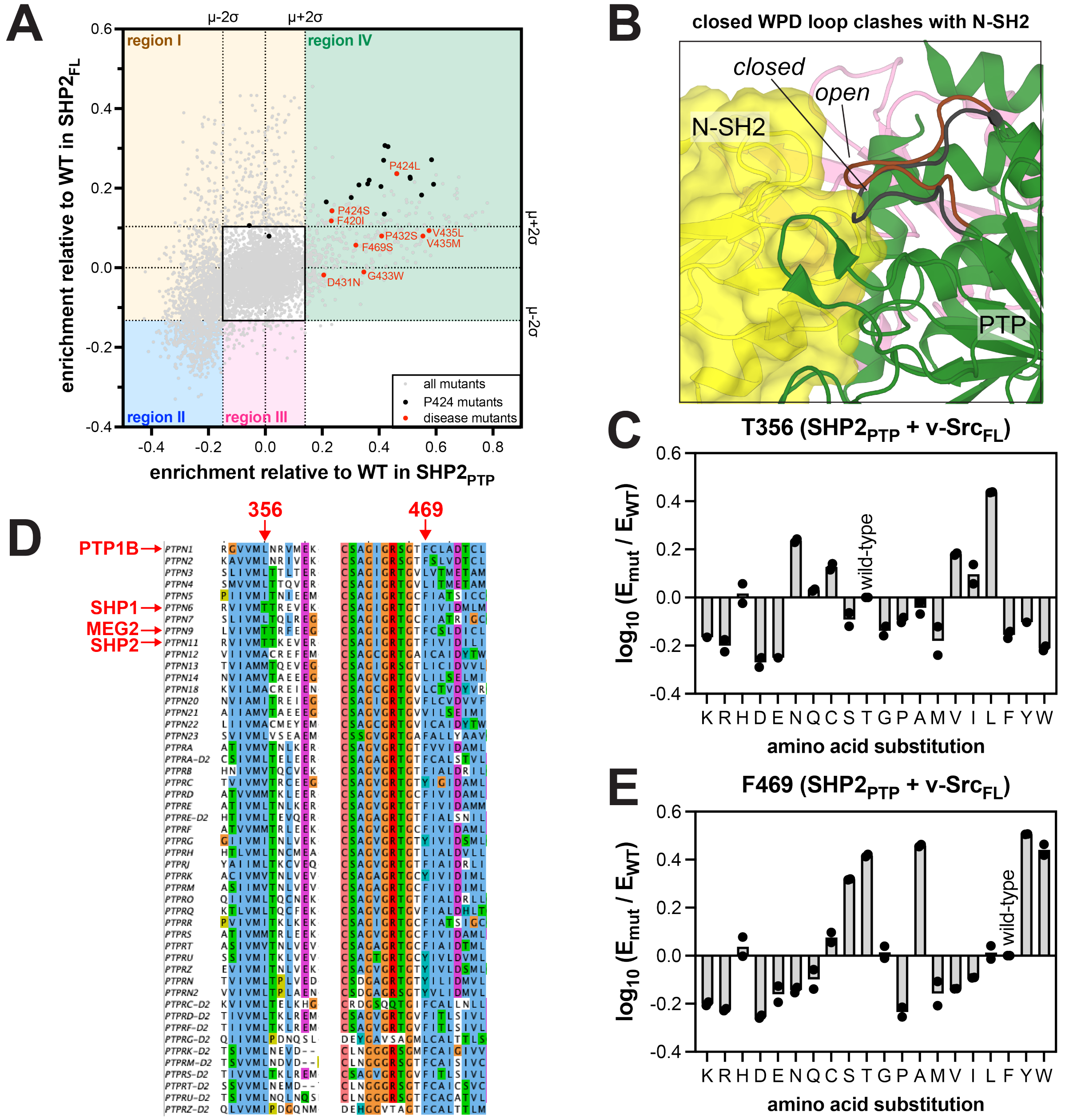


**Supplementary Figure 5. PTP domain mutations that modulate WPD loop structure and dynamics.** (**A**) Scatterplot juxtaposing SHP2$_{FL}$ and SHP2$_{PTP}$ mutational scanning datasets, highlighting the high density of P424 mutations in region IV, along with several unstudied disease mutants near the WPD loop. (**B**) The closed conformation of the SHP2 WPD loop (black) clashes with the N-SH2 domain (shown in yellow) in the autoinhibited state. The open conformation of the WPD loop (brown) does not clash with the N-SH2 domain. (**C**) Mutational effects at T356 in the SHP2$_{PTP}$ selection with v-Src$_{FL}$. (**D**) Alignment of human classical PTPs in the regions around SHP2 T356 and F469. (**E**) Mutational effects at F469 in the SHP2$_{PTP}$ selection with v-Src$_{FL}$.

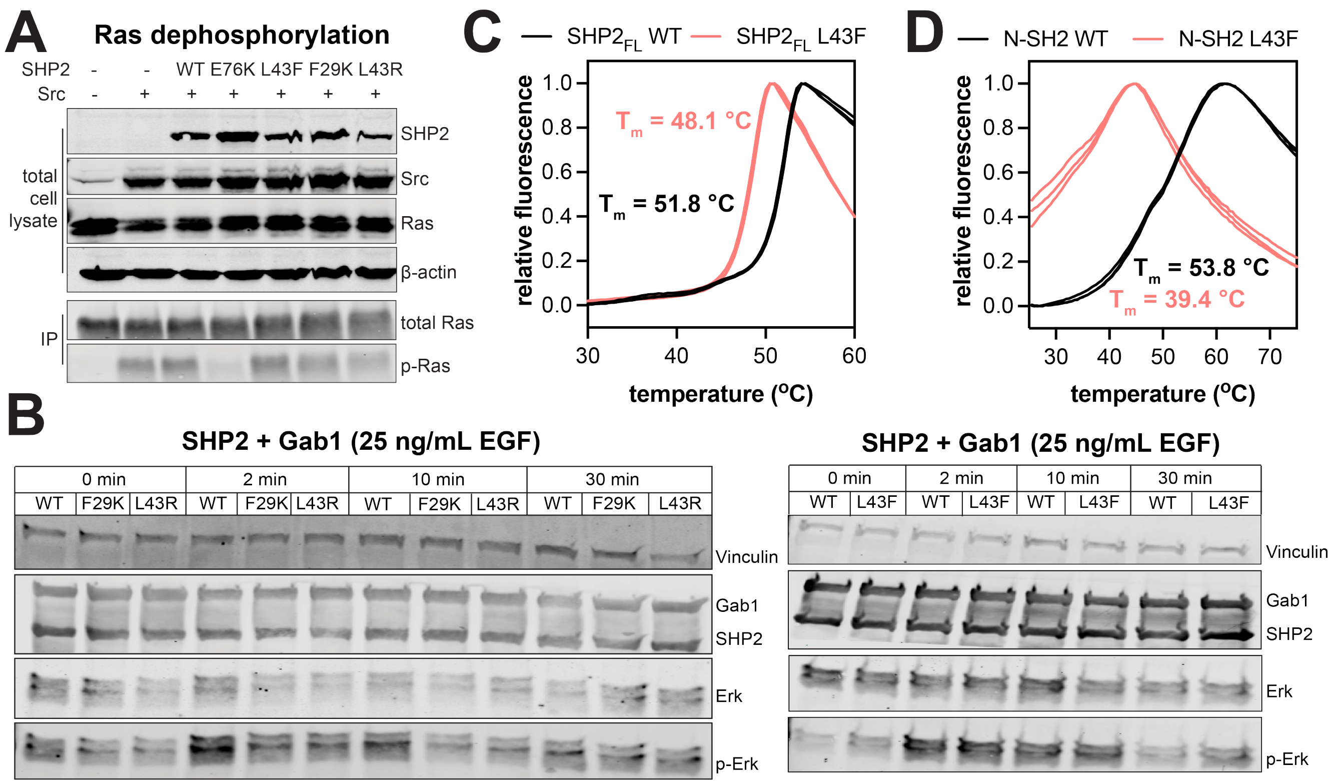


**Supplementary Figure 6. Destabilization of the N-SH2 domain by core mutations alters SHP2 activity.** (**A**) Representative western blot showing Ras dephosphorylation by SHP2 variants in HEK293 cells. (**B**) Representative western blot of SHP2 variant-dependent phospho-Erk levels upon EGF stimulation of HEK293 cells. (**C**) Differential scanning fluorimetry of the wild-type and L43F mutant $SHP2_{FL}$. (**D**) Differential scanning fluorimetry of the wild-type and L43F mutant isolated N-SH2 domains.

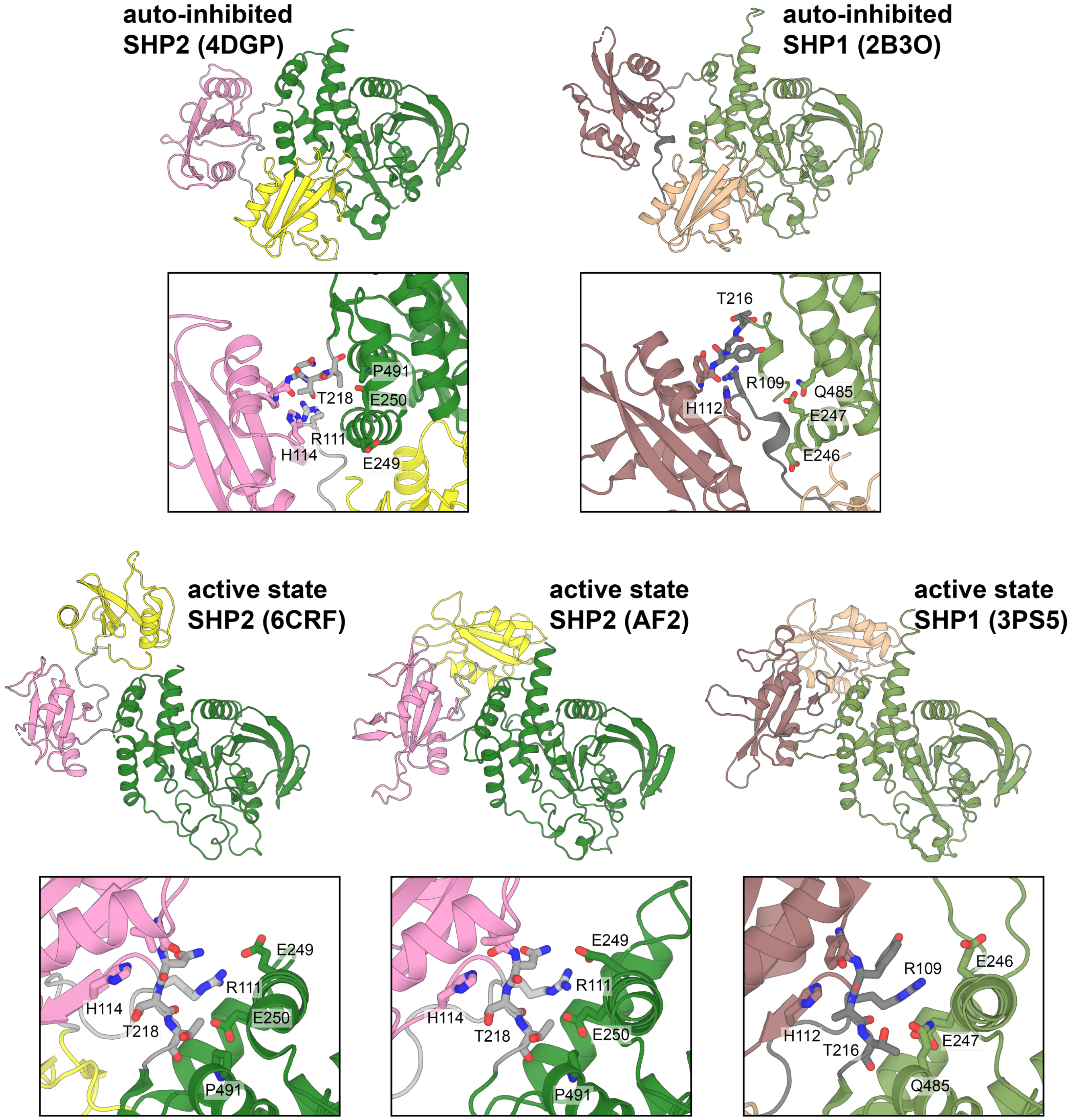


**Supplementary Figure 7. Divergent C-SH2/PTP interface in SHP1 and SHP2.** Structures of SHP2 and SHP1 in the auto-inhibited and active states are shown. Full-view structure images are aligned on the PTP domains (green) and highlight differences in C-SH2 orientation relative to the PTP domain across the models. The zoomed-in panels show similar views of the C-SH2/PTP interface in auto-inhibited state, with select residues highlighted. A different view is shown for the active state structures, with the same key interface residues highlighted.
